# High diversity of pimpline parasitoid wasps (Hymenoptera, Ichneumonidae, Pimplinae) from the lowermost Eocene Fur Formation

**DOI:** 10.1101/2021.11.18.468631

**Authors:** Seraina Klopfstein

## Abstract

With an estimated 100,000 extant species, Darwin wasps (Ichneumonidae) are more specious than all vertebrates together. However, only 288 fossil species have been described to date, with hundreds more awaiting formal description in palaeontological collections. One of the largest gaps in our knowledge concerns the ~12 million years between the K-PG mass extinction and the late Ypresian, from which only two species have been formally described, including *Pimpla stigmatica* Henriksen from the Danish Fur Formation (~55 Ma). Ie here redescribe and reclassify this species in the genus *Epitheronia* Gupta, n. comb., and describe nine new species from this fossil locality that are consistent with a placement in Pimplinae: *Crusopimpla collina* n. sp., *C. elongata* n. sp., *C. minuta* n. sp., *C. rettigi* n. sp., *C. violina* n. sp., *Theronia? furensis* n. sp., *T. nigriscutum* n. sp., *Xanthopimpla ciboisae* n. sp., and *X. crescendae* n. sp. The diagnosis of the genus *Crusopimpla* Kopylov, Spasojevic & Klopfstein is amend in the light of the new species. By comparing the preserved colouration between and within specimens of different species, we draw conclusions about the taxonomic usefulness of colour patterns observed in Fur Formation ichneumonids. The number of described species of pimpline parasitoid wasps from Fur is very high when compared to any other fossil deposit, but low with respect to numbers of extant species. Further study and excavation of Fur ichneumonids will certainly reveal an even higher diversity.

## Introduction

The fossil record of Darwin wasps (Hymenoptera, Ichneumonidae; Klopfstein et al. 2019b) has received even less attention than their extant diversity, with only 288 described fossil species facing more than 25,000 recent ones (Yu et al. 2016). As a consequence, our understanding of the evolution of this species-rich taxon through time is very limited, hindering inferences about the structure of past ecosystems in which these parasitoid wasps presumably played a similarly important role as they do today. The family is rather easy to identify even as fossils due to their unique and remarkably constant wing venation pattern (Broad et al. 2018): even isolated fore wings of can be identified robustly as belonging to this family. And much like in many terrestrial habitats today, numbers of both individuals and species of this group are high in many Cenozoic and even some Mesozoic deposits (Brues 1910, Kopylov 2009, 2010, Kopylov et al. 2018, Spasojevic et al. 2018b). In contrast to family placement, classifying fossil Darwin wasps into one of the 40 extant and four extinct subfamilies is typically very difficult; the obstacles adherent to the interpretation of fossil insects in general are exacerbated by high levels of homoplasy in this family caused by parallel adaptations to the same host groups (Gauld & Mound 1982). Most subfamilies can only be diagnosed based on a combination of multiple characters, and fossil placement thus often remains highly ambiguous, except in very well-preserved fossils (Klopfstein & Spasojevic 2019).

The first appearance of ichneumonids in the fossil record dates back to the lower Cretaceous or even upper Jurassic (Kopylov 2009, Zhang & Rasnitsyn 2003), but only a single specimen from the Mesozoic is currently classified in an extant subfamily: *Albertocryptus dossenus* Mckellar, Kopylov & Engel from Canadian amber (Mckellar et al. 2013). The remaining 61 Darwin wasps from the Mesozoic were described in four extinct subfamilies, none of which has been recorded from the Cenozoic. The fossil record thus implies a strong taxonomic turnover of Darwin wasps, as it could have resulted from the K-Pg mass extinction event. However, this hypothesis is on rather shaky grounds for several reasons. First, the phylogenetic affinities of the Mesozoic subfamilies remain poorly known, and they could in fact represent early lineages of extant subfamilies, instead of truly extinct radiations (Kopylov 2009, 2010). For instance, some genera currently classified in the extinct Labenopimplinae, such as *Rugopimpla* Kopylov, might well represent stem representatives of extant subfamilies (D. Kopylov, personal communication). Second, a recent molecular study (*Spasojevic et* al. 2021) has estimated the age of the radiation of the nine subfamilies that form the clade called Pimpliformes to the Middle Jurassic period, implying the appearance of most extant subfamilies deep in the Mesozoic. Even though such an old age estimate for Pimpliformes is at odds with the oldest known fossil attributed to this group, which stems from the late Paleocene (Piton 1940), the size of the gaps in the described fossil record of Darwin wasps implies that it is far from unlikely. While gaps in the fossil record are common even in well-studied insect groups, their length is almost certainly further increased in Darwin wasps by the neglect of this group by palaeoentomologists.

The best-studied epoch for Ichneumonidae fossils is clearly the Eocene, with 146 fossil species described, mostly from five famous insect fossil localities: Messel pit (Spasojevic et al. 2018b), Green River (Scudder 1890, Spasojevic et al. 2018a), Baltic Amber (Kasparyan 1988, Kasparyan & Humala 1995), Isle of Wight (Antropov et al. 2014), and Florissant shales (Brues 1910). In contrast, only two species have been described from Cenozoic strata that are older than these: *Phaenolobus arvernus* Piton from the early Paleocene Menat Formation in France (Piton 1940) and *Pimpla stigmatica* from the early Eocene Fur Formation in Denmark (Henriksen 1922). In an attempt to improve our knowledge of fossil Darwin wasps, we studied the latter formation in more detail.

The Fur Formation is a marine lagerstätte deposited right after the Paleocene-Eocene Thermal Maximum (PETM) greenhouse event, which ended about 55 Ma (Westerhold et al. 2009). These deposits are rich in marine taxa, but terrestrial groups can also be found, including some of the oldest representatives of modern birds and more than 20,000 insect specimens, most of which were gathered by private collectors. Both preservation and sheer abundance of insects from Fur are outstanding (Henriksen 1922, Larsson 1975, Rust 1998, 2000). The deposit has already revealed conclusive evidence for a moth mass migration (Rust 2000), and the exceptional preservation of the complex eyes of tipulid flies allowed drawing parallels to the micromorphology of trilobite eyes (Lindgren *et al*. 2019). The large number of fossils even from groups such as Diptera, which fossilize poorly compared to more strongly sclerotized insects like beetles, might allow reconstruction of past insect communities; however, most groups still await detailed taxonomic study (but see Kohring 1994, Rust & Møller Andersen 1999).

Darwin wasps from the Fur Formation have already received some attention in a taphonomic treaty of the insects found in these deposits (Rust 1998), and their appearance in a marine fossil record, about 100 km from the nearest coast, has been discussed in relation to observations of migratory behaviour in extant species across the open sea (Ansorge 1993, Horstmann 1970). However, only a single Darwin wasp species from Fur has been formally described (Henriksen 1922). Rust (1998) pointed out the existence of two distinct colour morphs among the material he examined and linked them to differences in the predominant body position they had fossilized in. Beside these two morphs, additional patterns of colouration are evident among the Darwin wasp specimens that have accumulated since, but the interpretation of these patterns require some careful consideration. In fossils, colouration is typically altered strongly during diagenesis (Mcnamara 2013). Colours and even colour patterns in fossils thus cannot be taken at face value when interpreted in a taxonomic context. Nevertheless, the often very detailed patterning seen in many insect fossils might still reflect at least part of the original colour pattern, an assertion that has recently gained momentum by the discovery of preserved melanin pigments in fossils, including in a fish eye from the Fur Formation (Lindgren *et al*. 2012).

We studied more than 110 ichneumonid specimens from the Fur formation and found an astonishing diversity (Fig. 1), with several extant subfamilies evidenced by multiple species. Here, we focus on those specimens that show characters that are consistent with a placement in the extant subfamily Pimplinae. Members of Pimplinae cannot be diagnosed based on a single character, but rather on a combination of several characters, with some of the most decisive ones only rarely visible in fossils (but see Spasojevic et al. 2018b). Nevertheless, we found and describe nine new species with robust character evidence for a placement in Pimplinae. Furthermore, we redescribe *Pimpla stigmatica* and revise its generic placement in the light of additional specimens of this taxon. Finally, we examined patterns of coloration present in the Fur Formation pimplines, comparing them both within and between the studied pimpline specimens, as well as to their extant counterparts, to evaluate their usefulness for species delimitation and classification.

**Figure 1.**
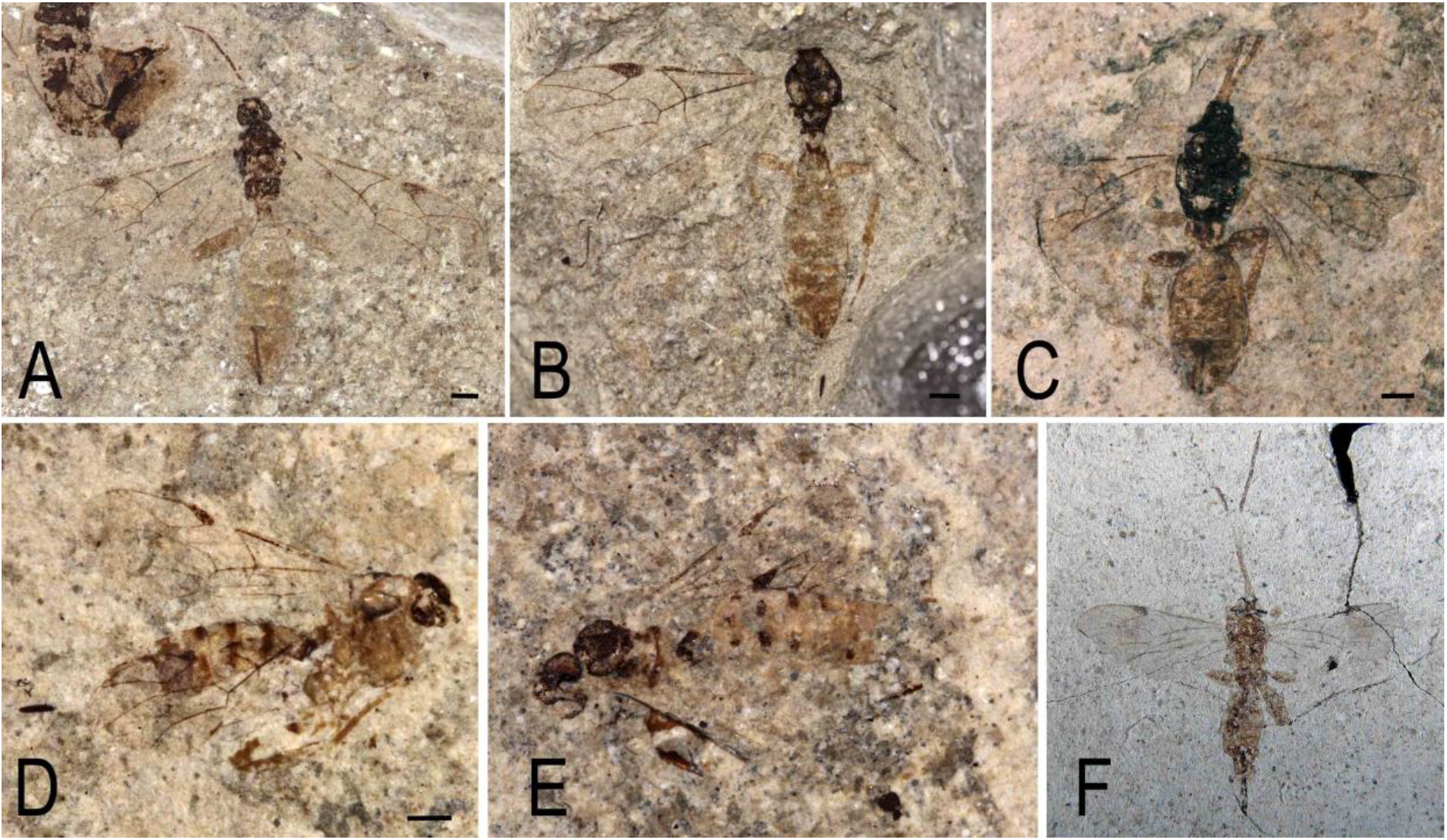
Examples of colour preservation in pimpline Darwin wasps from the Fur Formation. A. Paratype #13817 of *Crusopimpla violina*. B. Paratype #11216 of *C. violina*. C. Paratype #11219 of *C. rettigi*. D. Paratype #13065 of *Xanthopimpla crescendae*. E. Paratype #10918 of *X. crescendae*. F. Specimen #12707 of *Epitheronia stigmatica*.

## Material and Methods

All the fossils described herein stem from the Fur Formation in Denmark, a marine sediment deposited during the Ypresian (Lower Eocene). They were collected mostly by the private collectors Erwin Rettig, Jan Verkleij and Ole Burholt and by the former director of the Fur Museum, Magne Breiner Jensen. They are kept at the Fur Museum in Nederby (FUR) and the Fossil- and Mo-clay Museum in Nykøbing Mors (MOL).

All fossils were studied under a Leica WILD M8 stereo microscope and photographed using a Leica M205C stereomicroscope and the Imagic IMS Client software. Interpretative drawings were created in Adobe Photoshop CS5 using separate layers for each body part and for the part and counterpart of a fossil, if available. In the drawings, solid lines represent rather certain and dotted lines uncertain interpretations. In some cases, specific colour patterns are indicated by dotted areas. For the interpretation of colouration, both part and counterpart of a fossil and the colour and patterning of the surrounding rock were considered.

Morphological terminology follows Broad et al. (2018), with numbers for the respective sections added to wing veins as in Figure 2. To denote tergites and sternites, we used the abbreviations T1, T2 and S1, S2 and so on. Measurements were taken in Image J. Unless stated otherwise, measurements correspond to the longest length of a structure. When several specimens were included in a species, measurement ranges are given, with the values of the holotype repeated in brackets. If start or end point of a measurement had to be guessed, then the measurement is preceded by a tilde sign (“~”). The length of the often fragmented antennae is given both as the minimum length based on clearly visible segments and the estimated (i.e., interpolated) length, if they are different. Fore wing and body length (from head to last tergite, excluding the ovipositor) are given as absolute values, while the remaining measurements are taken as ratios, with forewing length as the denominator except if indicated otherwise. The length of the pterostigma is measured from the end of the costal vein. The length of the ovipositor is measured, as is custom in Darwin wasps, from the attachment point of the ovipositor sheaths to the tip.

**Figure 2.**
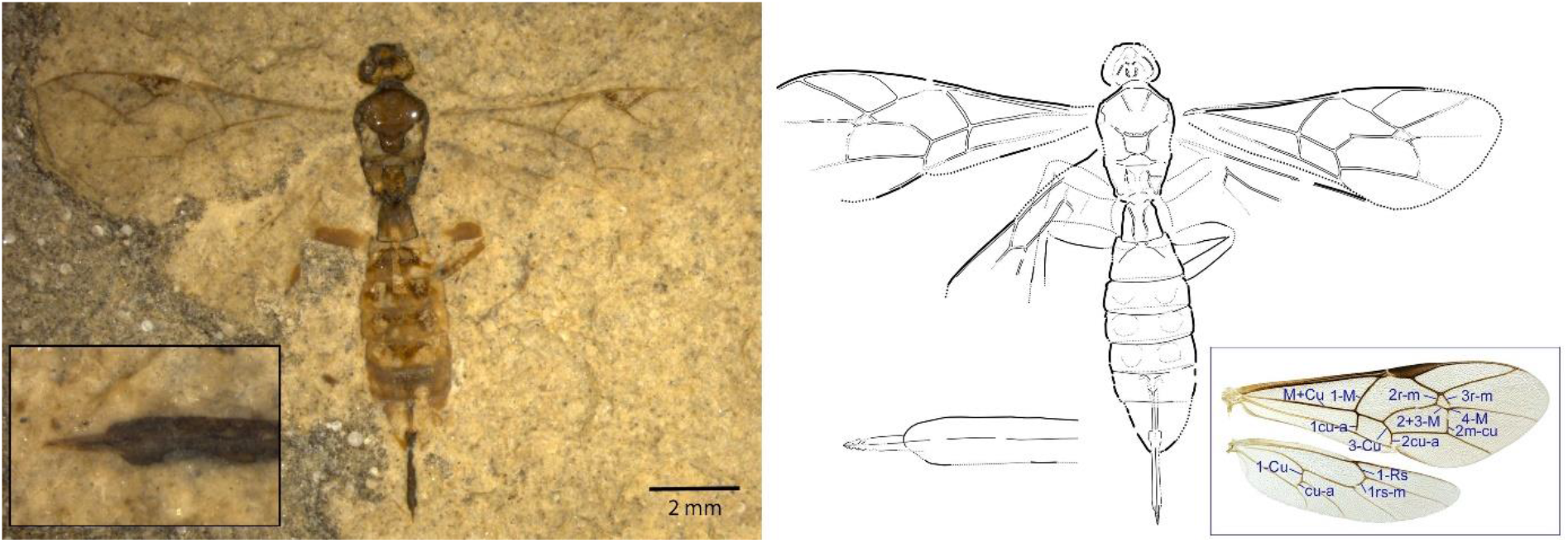
Holotype of *Crusopimpla collina* (#14680), photograph and interpretative drawing, where dotted lines represent uncertain and/or interpolated interpretations. The blue box on the lower right shows wing vein names on photographs of the extant Darwin wasp *Rhimphoctona obscuripes* (Holmgren, 1860).

Species delimitation in fossils can be tricky, as incomplete preservation might not allow comparison of all relevant characters and thus leave considerable uncertainty about the association of different fossils. We followed a cautious approach when denoting new species, which we only considered as distinct if several characters show clear differences. Many fossil specimens were too incomplete for them to be associated firmly with a species; we placed them tentatively within a species, but they do not form part of the type series and were not considered for the species descriptions.

## Results

### Subfamily identification

We used the following character states to diagnose members of Pimplinae (Table 1): a) forewing with areolet triangular or oblique-quadrate (rarely open); b) forewing vein 2m-cu bowed outwards, with two bullae; c) first tergite attached broadly to propodeum, parallel-sided or somewhat converging towards the base; usually 1.0-2.0 times longer than wide, sometimes longer; d) spiracle of first tergite in front of the middle; first sternite shorter than half the length of its tergite; e) ovipositor at least as long as apical depth of metasoma, usually longer; if shorter than metasoma, then usually rather robust; f) ventral valve of ovipositor with teeth, dorsal valve without a notch; g) claws usually with a basal lobe. We here considered a fossil species as belonging to Pimplinae if it showed at least three of the above characters and did not clearly contradict any of the others. Following these criteria, we found about 74 specimens that could belong to this subfamily. Out of these, we delimited ten pimpline species (Table 1), represented by 25 specimens that could be placed with some certainty in these species, plus another 22 with a tentative placement in one of them. The remaining specimens were too poorly preserved to identify them any further at this stage.

**Table 1.**
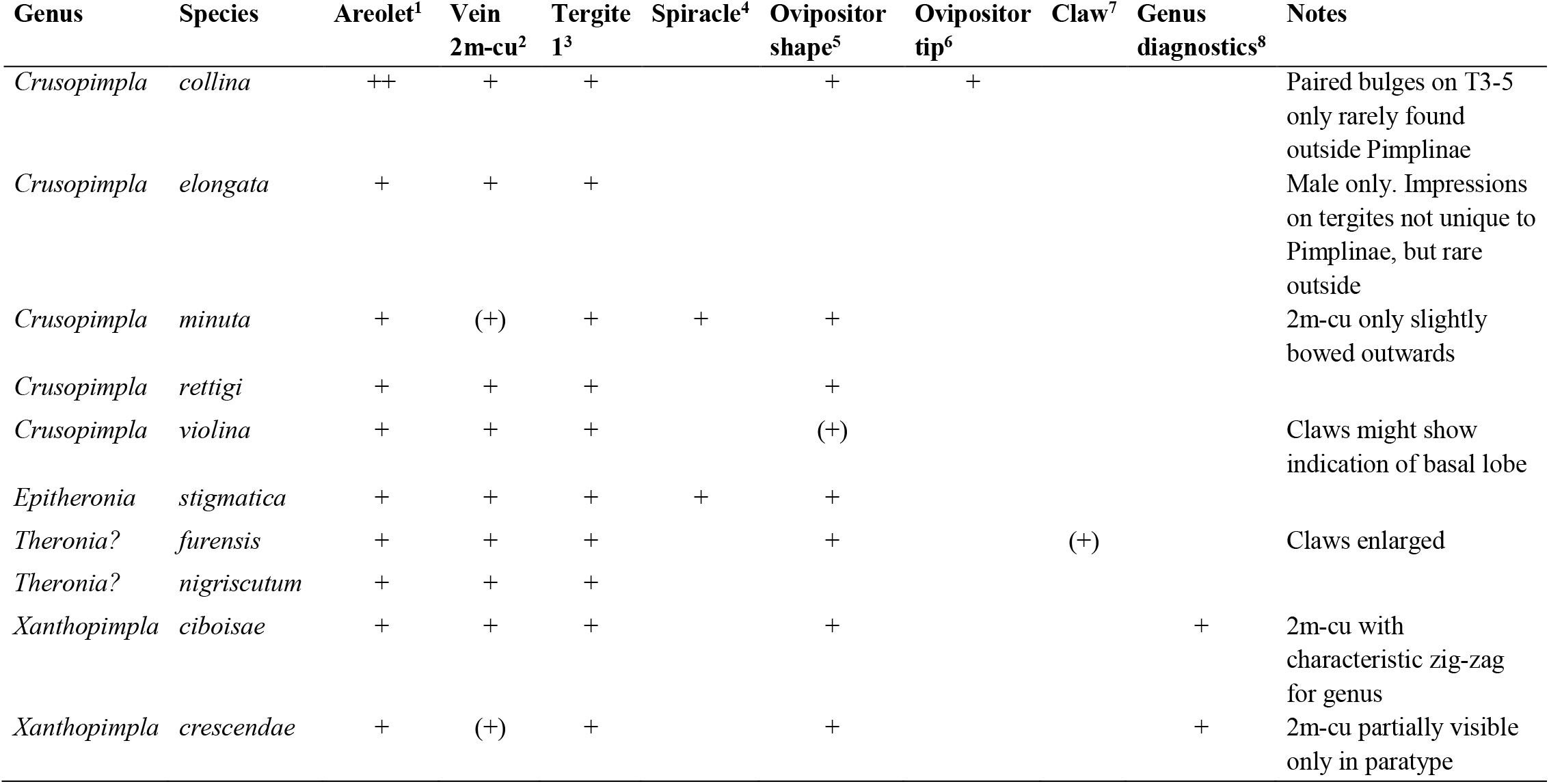
Overview of characters supporting placement in Pimplinae. Brackets indicate uncertain interpretation or visibility of only some aspect of a character. 1, Forewig with areolet triangular or oblique-quadrate. 2, Forewing vein 2m-cu bowed outwards, with two bullae (in brackets if bullae not visible). 3, First tergite attached broadly to propodeum, parallel-sided or somewhat converging towards base; usually 1.0-2.0 times longer than wide. 4, Spiracle of first tergite in front of middle; first sternite shorter than half the length of tergite. 5, Ovipositor clearly protruding from metasomal apex, often long; if shorter than metasoma, then usually rather robust. 6, Ventral valve of ovipositor with teeth, dorsal valve without a notch. 7, Claws with basal lobe (present in many extant Pimplinae, especially Ephialtini). 8, Unique character combination present that allows association with a particular genus in Pimplinae.

### Preservation of colours and colour patterns

We found rather heterogeneous preservation of colours and colour patterns across the studied pimpline specimens from the Fur formation (Fig. 1; see also images of holotypes under the mentioned species). On the one hand, all type specimens of *Crusopimpla rettigi, C. violina* and *Theronia? furensis* showed very consistent colouration, both across and within specimen, which can be assumed to correspond rather well with their original colour pattern, even if the actual colouration might be somewhat altered. All these fossil species share a dark brown head and mesosoma and orange metasoma and legs (Figs 1A-C). This colour pattern is still very common in the family and also occurs in several genera of Pimplinae.

In contrast, some of the more lightly coloured fossil specimens exhibit inconsistencies in colour patterns, which manifest either between specimens (for instance between the holotype and additional specimens of *Epitheronia stigmatica*) or even within a single specimen (Figs 1C-F). The latter is evident from an apparent asymmetry for instance in dark paired patches on the tergites of some specimens of *Xanthopimpla* Saussure (Fig. 1D), or from the irregular colouration of the mesoscutum of specimens of *Xanthopimpla* and *E. stigmatica*, which in some cases retained its originally dark colouration only along the outer margin. Without several specimens of the same species at hand, irregular preservation of colour can lead to misinterpretations. For example, the antenna of *E. stigmatica* (Fig. 1F) has an orange base, light middle and dark brown tip, but without comparing several specimens that consistently show this pattern, the light middle part of the antennae might easily have been interpreted simply as being less-well preserved than the rest, instead of reflecting a truly lighter colouration.

Overall, it appears that dark colouration is more consistently preserved than light colouration, but this seems to depend also on the ground colour of a species: the dark colouration present in some of the more light-bodied species appears far less consistently preserved than the same body parts in specimens with already dark ground colour. In addition, where general preservation differs strongly between specimens, preserved colouration often also does, like in the only specimen of *Crusopimpla collina*: along with its nearly three-dimensional preservation, it shows such irregular colouration that it is not possible to reconstruct its original colour pattern without access to additional specimens of this species. Bearing these difficulties in interpretation in mind, we here describe colouration as found in the fossils, but integrate over all available specimens for taxonomic interpretation. For species delimitation, we rather focus on colour patterns than on absolute hues.

### Key to species of the Fur Formation

This key includes the species currently known from the Fur Formation, but should be expanded as soon as additional species are being described.

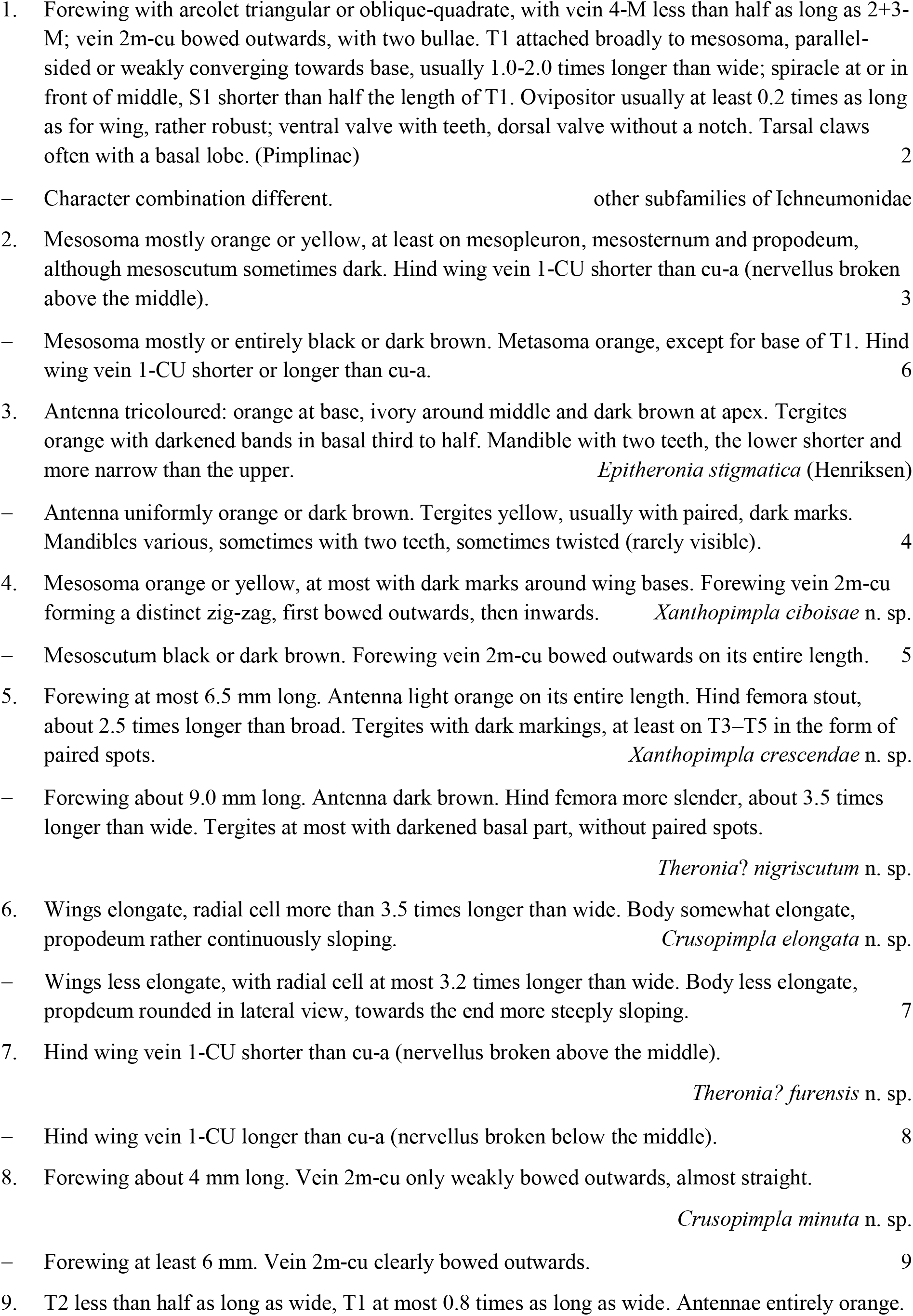
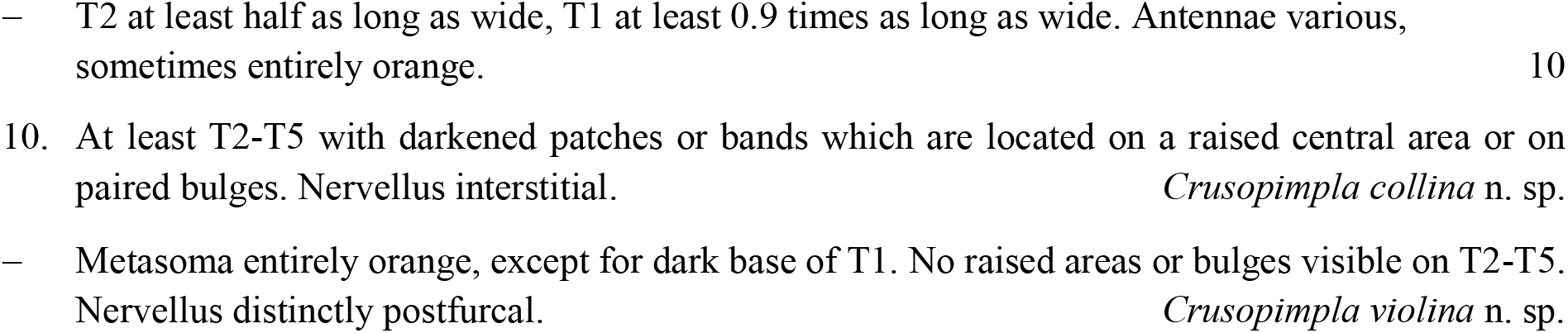

### Systematic Palaeontology

*Crusopimpla* Kopylov, Spasojevic & Klopfstein 2018

*Crusopimpla collina* **n. sp.**

Figure 2

**Type material**. Holotype female: #14680 (part and counterpart; FUR; leg. H. Breiner).

**Etymology**. This fossil is unusual in its three-dimensional preservation, which allows distinguishing not only the structure of the mesosoma in great detail, but also reveals paired bulges present on T3-T5. It owes its name to these little hills on the metasoma.

**Type horizon and locality**. Denmark, Jutland, Limfjord region, Skive kommune, Fur, cement stone.

**Diagnosis.**

*Taxonomic placement*. Most of the characters that we use here to diagnose Pimplinae are well visible in this specimen (Table 1), and it in terms of body proportions would fit well among some smaller-sized ephialtine genera such as *Scambus* Hartig, *Gregopimpla* Momoi or *Iseropus* Förster. However, the exceptionally well-preserved propodeum reveals a nearly complete carination, which is not a character that is seen anywhere in the subfamily today. As different tribes and genus groups within Pimplinae feature different carinae being reduced, Kopylov et al. (2018) concluded that a full set of carinae is probably the plesiomorphic state in the subfamily. Based on a fossil from the Eocene Tadushi formation, they described a new genus, *Crusopimpla*, which shows extensive propodeal carination. The two species currently placed in *Crusopimpla* (*C. tadushiensis* Kopylov, Spasojevic & Klopfstein and *C. redivia* (Brues)) only have the forewings and not the hindwings preserved, which is why any statement about the relative lengths of veins 1-Cu and cu-a (which together form the nervellus) is missing from the original description of this genus. Our species clearly has the nervellus broken below the middle, with vein 1-Cu about 1.5 times as long as cu-a. It is unclear what the plesiomorphic state of this character might be in the subfamily, but we currently certainly cannot use this character to rule out placement in this genus. We thus place our fossil here and thus expand the generic definition by this character. Also, the genotype *C. tadushiensis* appears to be dark coloured, although that might be an artefact of preservation, while *C. redivia* and the specimens added to the genus here show light brown or orange metasomas.

*Species diagnosis*. Compared to the genotype *Crusopimpla tadushiensis*, *C. collina* has more slender tergites (e.g., T2 0.55 times as long as wide, versus 0.32 in *C. tadushiensis). Crusopimpla? rediviva* is preserved only in lateral view, making measuring of tergite proportions difficult. But the latter species has a much more slender pterostigma (4.2 times longer than wide, versus 3 times in the current species). The two species previously in the genus also do not show any convex bulges on the tergites, even though that characters might have been obscured during fossilization.

**Description.**

*Preservation*. Body in dorsal view. Head probably turned around on the neck, thus showing from below and from the back; antennae missing. Mesosoma rather well preserved, including propodeal carination and details of the scutellar and postscutellar region; fore wings both nearly complete, partial hind wings present; partial legs visible. Metasoma almost complete, including sculptural elements on tergites; ovipositor sheaths well preserved, tip of ovipositor exposed and showing minute details.

Body 9.1 mm. Head and mesosoma dark brown; wing veins brown, pterostigma brown except for narrowly light area at base; legs orange. Metasoma orange, dark brown on T1 and with dark brown patches of various shapes on the remaining tergites (although that might be artefactual); ovipositor orange, its sheaths black.

*Head* with orientation somewhat uncertain: clearly showing hind side of head, with distinctive foramen magnum and genal carina, but might have turned around on its stalk and showing the ventral side.

*Mesosoma* rather short and stout; deep notauli converging on basal 1/3, then no longer visible; tegulae indicated by lighter areas; prescutellar groove with deep pits on either side, scutellum rather short, and convex; postscutellum outlined; propodeum with distinct carination, corresponding to nearly complete lateromedian and at least partial lateral longitudinal and pleural carinae and at least partial basal and apical transverse carinae, including complete outline of right area externa and area dentipara; indication of either apophyses or extended hind corners of pleural carinae. *Fore wing* 7.0 mm; areolet closed, broad-quadrate to nearly pentagonal, with uneven sides, 4-M very short, 2r-m a bit shorter than 3r-m; 2m-cu with two bullae, bowed outwards on entire length; 1cu-a meeting M + Cu opposite 1-M; short ramulus present; 3-Cu longer than 2cu-a; radial cell 2.5 x longer than wide. *Hind wing w*ith 1-Rs about 1.7 x longer than 1rs-m, 1-Cu probably about 1.5 x as long as cu-a. *Mid and hind legs* not well preserved, but rather stout.

*Metasoma* with T1 about 0.95 x as long as wide, with strong median longitudinal carinae converging on basal half, then slightly diverging and finally converging again; T2 0.55 x times as long as wide, with basal oblique grooves cutting of anterolateral corners, seemingly convex in-between, colouration showing roundish patterns which probably indicate strong punctuation; T3 – T5 with paired roundish bulges that again show signs of strong punctuation; T7 clearly longer than T6. Ovipositor sheaths 0.22 x as long as fore wing; ovipositor tip extending from sheaths, with teeth-like outline clear at least on right side.

*Crusopimpla? elongata* **n. sp.**

Figure 3

**Figure 3.**
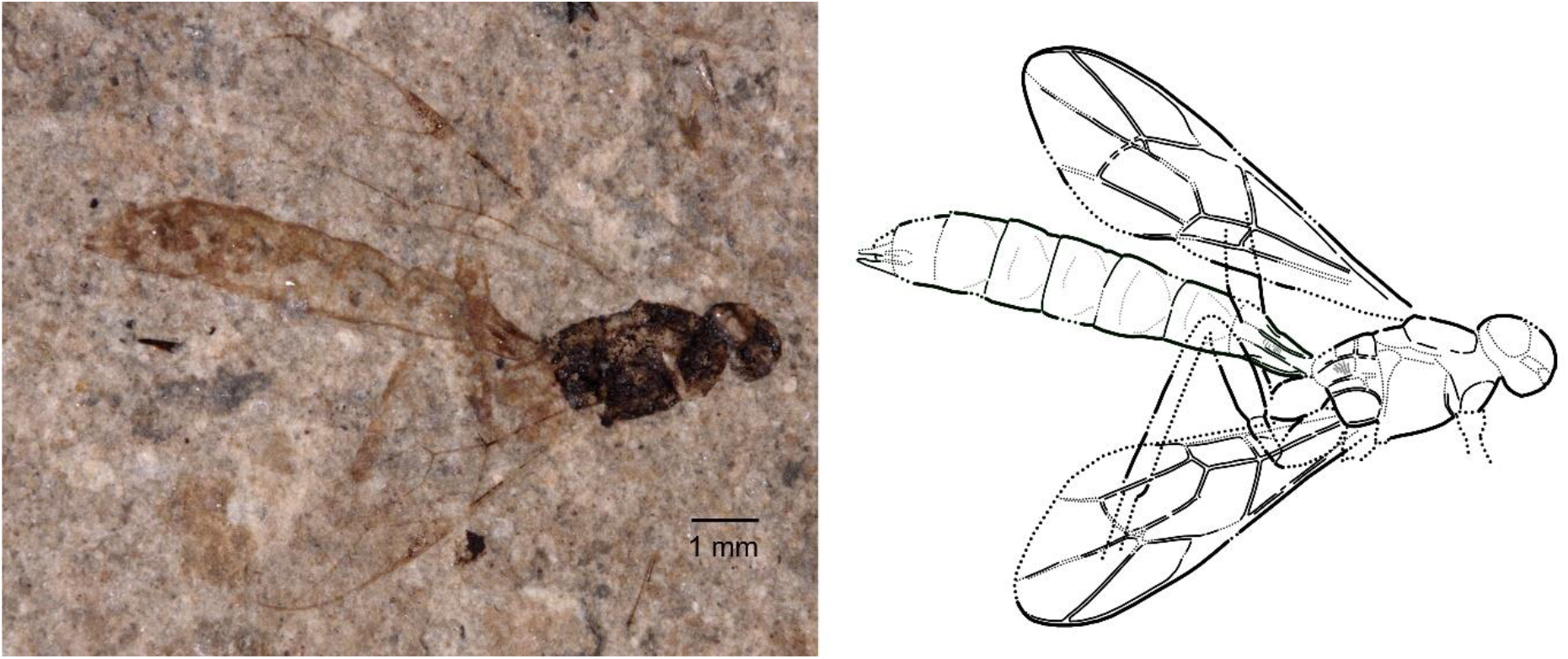
Holotype of *Crusopimpla elongata* (#11220), photograph and interpretative drawing, where dotted lines represent uncertain and/or interpolated interpretations.

**Type material**. Holotype male: #11220 (part and counterpart; FUR; leg. E. Rettig).

**Etymology**. The name of this species refers to its elongate body shape.

**Type horizon and locality**. Denmark, Jutland, Limfjord region, Morsø Kommune, Klinten ved Klitgård. Cement stone.

**Diagnosis.**

*Taxonomic placement*. Because only a single, though well-preserved male is available to base the description of this species on, subfamily placement is a bit less certain. However, the pattern of grooves and elevated areas on the tergites, which are showing rather consistently in both the part and the counterpart of this fossil, are rare outside this subfamily. It shares its colour pattern with *Theronia? furensis*, *Crusopimpla minuta, C. rettigi* and *C. violina* described here-in. The extensive carination of the propodeum points to the genera related to *Theronia* Holmgren or to *Crusopimpla*; placement in the latter is only tentative, as no hind wings are preserved and the state of the nervellus thus cannot be resolved.

*Species diagnosis*. The colour pattern is reminiscent of most other *Crusopimpla* described here and of *Theronia? furensis*. Furthermore *Crusopimpla collina* shows similarly elevated areas on its tergites. The current species differs from all of these and from the other species of the genus by its elongate wings and body.

**Description.**

*Preservation*. Holotype in lateral to latero-dorsal view. Head partially preserved, no antennae visible. Mesosoma rather well preserved, including part of propodeal carination; fore wings nearly complete, hind wings indiscernible. Metasoma highly complete, including well-preserved male genital organs.

Body 10.5 mm. Head and mesosoma dark brown, wing venation orange-brown, with light base of pterostigma. Legs orange. T1 brown at base, remainder orange, remaining tergites entirely orange; the very tips of parameres dark brown.

*Head* rather round, difficult to interpret; antennae missing.

*Mesosoma* a little elongate; notauli unclear; propodeum with distinct carination, corresponding to nearly complete pleural, lateromedian and maybe lateral longitudinal and partial transvers carinae, spiracle circular. *Fore wing* 7.1 mm; areolet closed, triangular, receiving vein 2m-cu at its outermost corner (4-M nearly obliterate); 2m-cu with two bullae, weakly bent outwards; pterostigma 4.4 x longer than wide; 1cu-a meeting M + Cu opposite 1-M. 1-Cu clearly longer than 2cu-a; radial cell 3.6 x longer than wide. *Hind legs* not well preserved, but rather slender.

*Metasoma* with T1 longer than wide, with strong median longitudinal carinae that are nearly parallel on most of its length; T2 at most a little wider than long, T3-T7 all nearly of the same length, with indications of diagonal grooves cutting off anterolateral corners and subapical transverse grooves. Parameres outlined, showing narrow and pointed tips.

*Crusopimpla minuta* **n. sp.**

Figure 4

**Figure 4.**
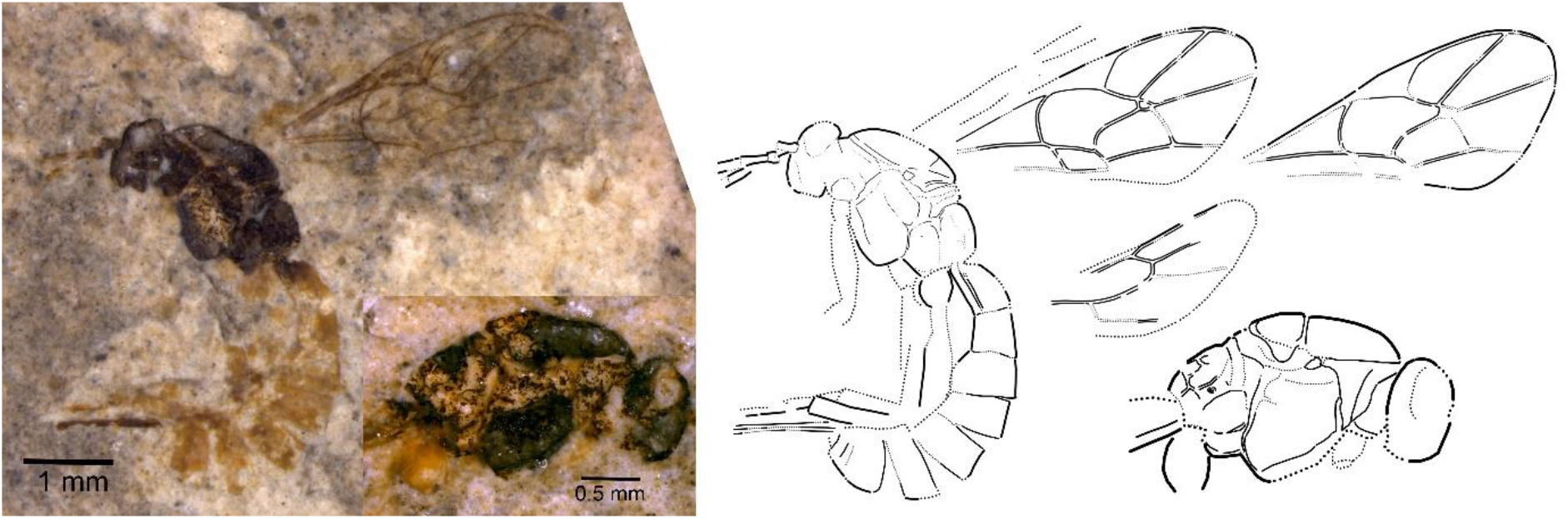
Holotype of *Crusopimpla minuta* (#13076), photograph and interpretative drawing, where dotted lines represent uncertain and/or interpolated interpretations.

**Type material**. Holotype female: #13076 (part and counterpart, FUR; leg. E. Rettig).

**Etymology**. The name of this species refers to its small size.

**Type horizon and locality**. Denmark, Jutland, Limfjord region, Morsø Kommune, Svalklit. Cement stone.

**Material examined**. Tentatively placed in this species: #14944 (FUR; leg. M. Breiner), #16636 (FUR; leg. O. Burholt).

**Diagnosis.**

*Taxonomic placement*. This species is very similar to the other species of *Crusopimpla* described herein. In terms of typical pimpline characters, the only somewhat equivocal one is the shape of vein 2m-cu, which is less outward-bowed in this species than in any other pimpline described her from the Fur Formation. However, this might just be a consequence of its small size. Placement in *Crusopimpla* then is not only supported by the very similar colour pattern to *C. rettigi* and *C. violina*, but also by the clearly visible propodeal carination. *Crusopimpla minuta* can be distinguished from any other species of the genus by its small size.

**Description.**

*Preservation*. Holotype in lateral view. Head and base of antennae preserved. Mesosoma rather well preserved, including details of metanotal area and part of propodeal carination; fore wings nearly complete, one partial hind wings partly discernible. Metasoma complete with ovipositor sheaths, the latter maybe broken off and only partial (unclear).

Body 5.3 mm. Head and mesosoma black or dark brown, wing venation orange-brown. Antennae entirely orange (holotype), as are the legs. T1 dark only at the very base, rest orange, remaining tergites entirely orange; ovipositor sheaths dark brown.

*Head* transverse, wider than long; only base of antenna preserved, with scape rather short and wide.

*Mesosoma* rather stout; notauli unclear, maybe indicated at front of mesoscutum; propodeum distinctly shortened, with nearly circular spiracle, pleural, lateral longitudinal and at least partial median longitudinal carinae present, both transverse carinae evident at least medially. *Fore wing* 3.8 mm; areolet closed, quadrate, with uneven sides, 4-M very short, 2r-m about as long as 3r-m; 2m-cu not well preserved, only very little bent outwards; pterostigma 2.7 x longer than wide; 1cu-a meeting M + Cu clearly distad from 1-M; 3-Cu clearly longer than 2cu-a; radial cell 2.4 x longer than wide. *Hind wing w*ith 1-Rs about 1.5 x longer than 1rs-m, 1-Cu maybe 1.5 x as long as cu-a. *Hind legs* not well preserved, but rather stout.

*Metasoma* with T1 showing an expansion just after its base in lateral view and an indication of what is probably the spiracle at about 0.25 its length; median longitudinal carina not well visible but maybe indicated; T2 probably shorter than wide (but lateral view), following tergites clearly transverse. Ovipositor sheaths at least 0.34 x as long as fore wing (tip not preserved), rather robust.

*Crusopimpla rettigi* **n. sp.**

Figure 5

**Figure 5.**
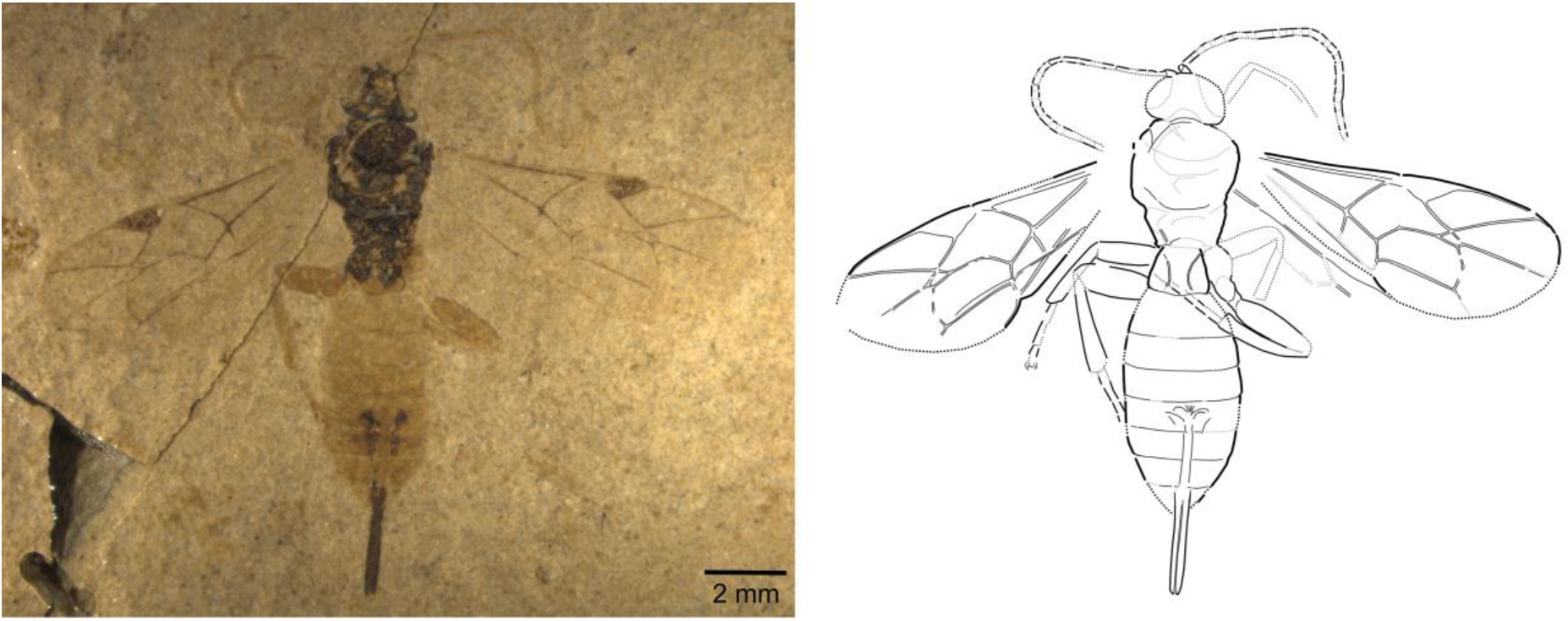
Holotype of *Crusopimpla rettigi* (#11209), photograph and interpretative drawing, where dotted lines represent uncertain and/or interpolated interpretations.

**Type material**. Holotype female: #11209 (part and counterpart; FUR; leg. E. Rettig). Paratypes (FUR; both females): #11219, #13074.

**Etymology**. This species is dedicated to Mr Erwin Rettig, who collected the three specimens included in the type series.

**Type horizon and locality**. Denmark, Jutland, Limfjord region, Morsø Kommune, Klinten ved Klitgård (holotype and #11219), Svalklit (#13074). Cement stone.

**Diagnosis.**

*Taxonomic placement*. This species is very similar to *Crusopimpla violina* – see reasoning for subfamily and genus placement there.

*Species diagnosis*. Together with *Crusopimpla violina*, *C. rettigi* can be distinguished from other species of the genus by their colour pattern. From *C. violina*, it can be distinguished by its much stouter tergites and the entirely orange antennae, while they are at least partly dark brown in the other species.

**Description.**

*Preservation*. Holotype and paratype #11219 in dorsal, paratype #13074 in lateral view. Head in rather well preserved, in #13074 including mandibles. Antennae nearly complete in holotype, only base preserved in paratypes. Mesosoma rather well preserved, including part of propodeal carination; fore wings in holotype complete, in paratypes partial, hind wings indicated but with carination only weakly preserved, a bit better in #11219. Metasoma in holotype complete, in #11219 missing most of ovipositor and in #13074 from T7 onwards.

Body 8.6–10.5 (10.5) mm. Head and mesosoma black or dark brown, wing venation dark brown in forewing, with light base of pterostigma, and light brownish in hind wings. Antennae entirely orange (holotype). T1 dark at base, orange at apex, remaining tergites entirely orange, legs orange; ovipositor sheaths dark brown.

*Head* transverse, wider than long; mandible with two teeth (#13074), the lower a bit shorter; antennae with maybe 24-30 segments, median flagellomeres only slightly longer than wide, apical ones quadrate.

*Mesosoma* rather stout; notauli only visible anteriorly, rest unclear; propodeum distinctly shortened, with some faint traces of carinae. *Fore wing* 6.1–8.25 (8.25) mm; areolet closed, quadrate, with uneven sides, 4-M very short, 2r-m about as long as 3r-m; 2m-cu with two bullae, bent outwards; pterostigma 2.4–2.6 x longer than wide; 1cu-a meeting M + Cu a little distad from 1-M; 3-Cu clearly longer than 2cu-a; radial cell 2.5–3.0 x longer than wide. *Hind wing w*ith 1-Rs about 1.9 x longer than 1rs-m, 1-Cu about 1.1 x as long as cu-a. *Hind legs* not well preserved, but rather stout, hind femur maybe 3.0–3.2 x longer than wide.

*Metasoma* with T1 about 0.7–0.85 (0.85) x longer than wide, with strong median longitudinal carina converging on basal half, then parallel and diverging towards the posterior end; T2 0.4–0.45 (0.45) x times as long as wide. Ovipositor sheaths 0.30 x as long as fore wing, rather robust.

*Crusopimpla violina* **n. sp.**

Figure 6

**Figure 6.**
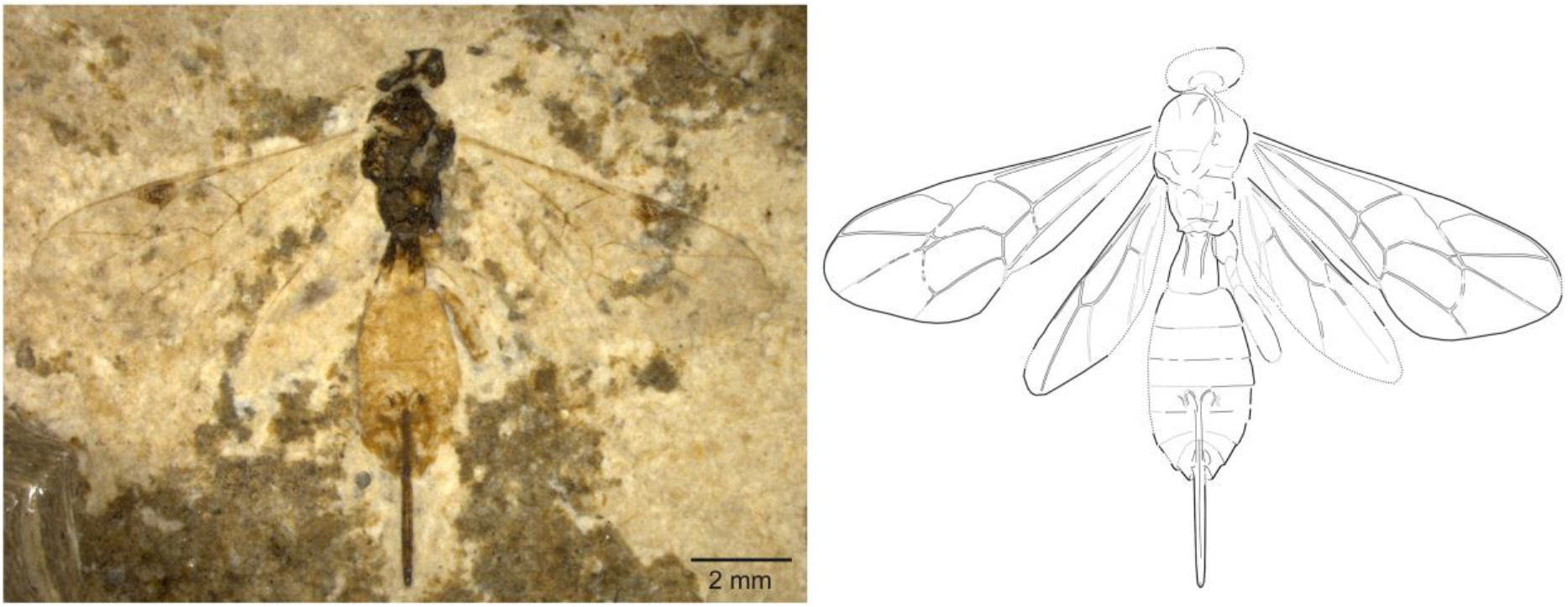
Holotype of *Crusopimpla violina* (#13061), photograph and interpretative drawing, where dotted lines represent uncertain and/or interpolated interpretations.

**Type material**. Holotype female: #13061 (part and counterpart; FUR; leg. E. Rettig). Paratypes (FUR; all females): #11212, #11269, #11216 (leg. E. Rettig), #13817, #16633 (leg. O. Burholt).

**Etymology**. This species is named for the elegant curves of the dorsomedian carinae on the first tergite, which remind of the shape of a violin.

**Type horizon and locality**. Denmark, Jutland, Limfjord region, Morsø Kommune, Svalklit (holotype), Klinten ved Klitgård (#11212, #11216, #11269), unknown (#13817, #16633). Cement stone.

**Material examined**. The following specimens are tentatively placed in this species: #10048, #10049, #10248, #11211, #11266, #11549, #13070, #16638.

**Diagnosis.**

*Taxonomic placement*. Subfamily placement of this species is somewhat less certain than in some of the other taxa (Table 1), as the ovipositor is somewhat more slender here. However, the remaining characters firmly point to this subfamily nevertheless. It shares its colour pattern (dark head and entire mesosoma, almost entirely orange metasoma and legs) with numerous extant ichneumonid species across many different subfamilies, including many Pimplinae, and with *Theronia? furensis*, *Crusopimpla minuta* and *C. rettigi* described here-in. From the *T. furensis*, it differs by the nervellus being broken clearly below the middle, a character not present in any species of the *Theronia* group nor the tribe Pimplini. The pronounced carination on the propodeum, even though not as clearly preserved as in *C. minuta* and *C. collina*, once more points to *Crusopimpla* as the most likely genus for this species, and it indeed shares many characters with *C. collina* as well as with the two species formerly included in this genus.

*Species diagnosis*. From the otherwise similar *C. rettigi*, this species can be distinguished by its longer tergites and at least partly dark brown antenna (paratype #13817) and from *C. minuta* by its much larger size. The colouration, especially the orange colouration of the metasoma distinguishes this species from all other *Crusopimpla* species.

**Description.**

*Preservation*. Holotype and paratypes all in dorsal view. Head in all cases partially preserved, in one paratype (#13817) including a few of the median flagellomeres. Mesosoma rather well preserved, including part of propodeal carination; fore wings in all four fossils complete or nearly complete, hind wings indicated but with carination only weakly preserved, somewhat difficult to interpret. Metasoma in all four types almost complete, in holotype and paratype #11269 including ovipositor sheaths and base of ovipositor.

Body 8.6–11.4 (8.6) mm. Head and mesosoma black or dark brown, wing venation dark brown in forewing, with light base of pterostigma, and light brownish in hind wings. Antennae dark brown dorsally, lighter ventrally. T1 dark at base, orange at apex, remaining tergites entirely orange, legs orange; ovipositor sheaths dark brown.

*Head* not well preserved, not much wider than long, antennae missing except in paratype #13817 which features some of the median flagellomeres, which are all at least twice as long as wide.

*Mesosoma* rather elongate compared to other pimplines; notauli converging on basal ¼, then parallel; scutellum of normal dimensions; propodeum with distinct carination, corresponding to nearly complete basal and probably also apical transverse, lateromedian and maybe lateral longitudinal and pleural carinae. *Fore wing* 7–8.8 (7) mm; areolet closed, quadrate, with uneven sides, 4-M very short, 2r-m a bit shorter than 3r-m; 2m-cu with two bullae, bent outwards; pterostigma 2.8–3.4 x longer than wide; 1cu-a meeting M + Cu opposite 1-M or a little distal from it (in holotype); 3-Cu clearly longer than 2cu-a; radial cell 2.5–3.1 x longer than wide. *Hind wing w*ith 1-Rs about 1.4 – 1.6 (1.5) x longer than 1rs-m, 1-Cu about 1.5 x as long as cu-a. *Hind legs* not well preserved, but rather stout.

*Metasoma* with T1 about 1.0–1.2 (1.2) x longer than wide, with strong median longitudinal carina converging at least on basal half, then diverging; T2 0.5–0.7 (0.7) x times as long as wide. Ovipositor sheaths 0.30–0.33 x as long as fore wing, rather robust.

*Epitheronia* Gupta, 1962

*Epitheronia stigmatica* (Henriksen, 1822) **n. comb.**

*Pimpla stigmatica* Hendriksen, 1822

Figure 7

**Figure 7.**
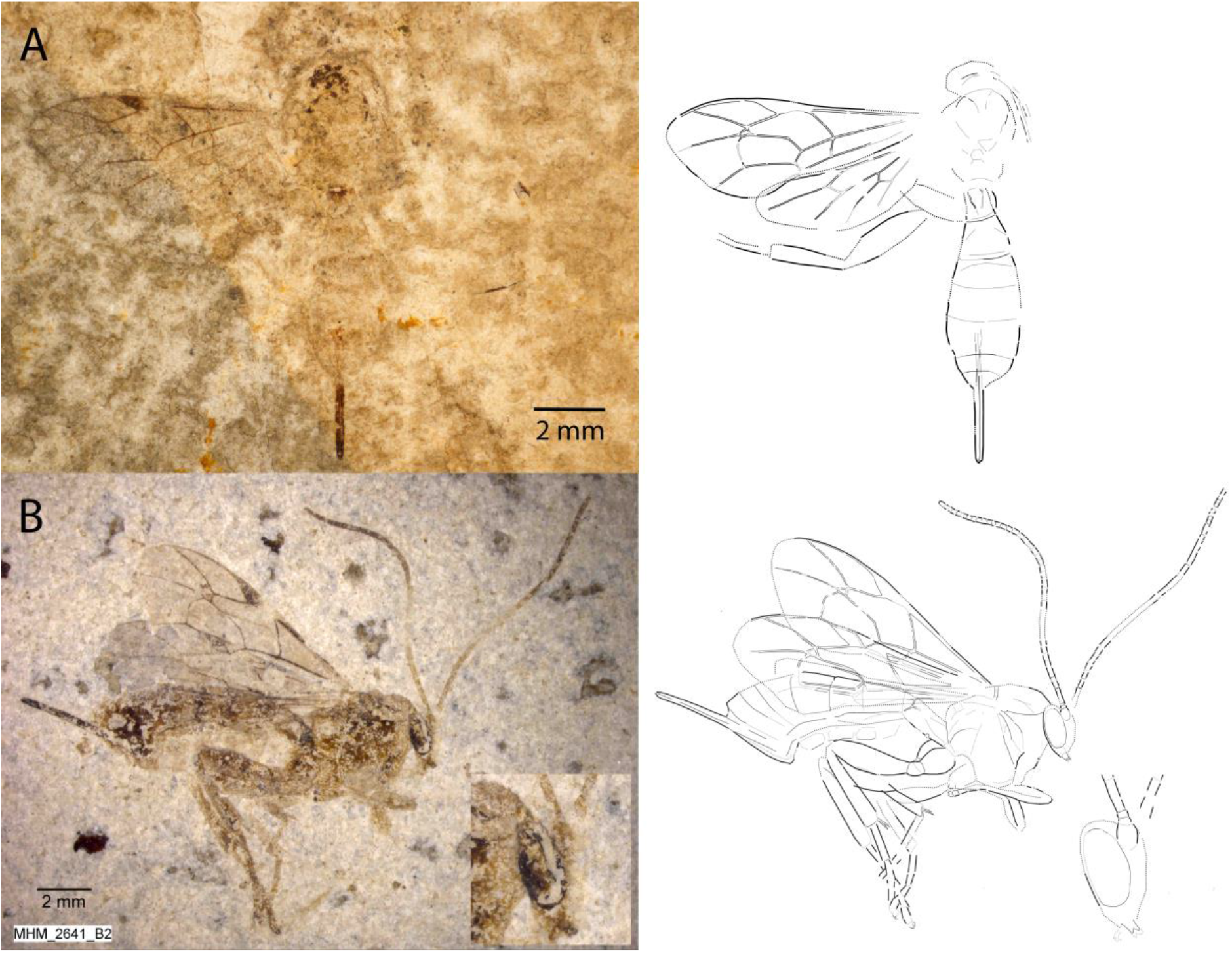
Specimens of *Epitheronia stigmatica*. A. Holotype, deposited at the Natural History Museum in Copenhagen. B. Specimen #MM-3141. Photographs and interpretative drawings, where dotted lines represent uncertain and/or interpolated interpretations.

The holotype of *Pimpla stigmatica* is rather poorly preserved, with the body mostly just outlined in the fossil, without showing any details. To avoid defining unnecessary new names, I here adopt this name for the species with the three-coloured antenna that is found rather commonly in the Fur Formation deposits. The venation of the fore and hind wing of the holotype fits rather well with these specimens, and there are no discernible characters on the body that would contradict the association, even though the holotype is at the lower end of the size spectrum.

**Type material**. Holotype examined at the Natural History Museum in Copenhagen.

**Type horizon and locality**. The type locality is given as “Thy” in original description, probably referring to the traditional Thy district in the Limfjord region in Jutland, Denmark. Additional specimens are from Denmark, Jutland, Limfjord region, Skive Kommune, Stolleklinten (#11922, #12707), Manhøje, Den nye grav (#11610); Morsø Kommune, Gullerup (#11110); and Ejerslev Molergrav, 56.918135°N, 8.918468°E (#MM-3141); unknown (#16635).

**Material examined**. All females: #MM-3141 (MOL), #10099, #11110, #11610, #11922, #12707, #16635 (all FUR). The following specimens are tentatively placed in this species: #9645, #10100, #11163, #11267, #11342, #13075, #15841.

**Diagnosis.**

*Taxonomic placement*. Henriksen correctly identified Pimplinae as the most likely subfamily. The areolet shape, outwards-bowed 2m-cu in the forewing, as well as the short and stout T1 and robust, protruding ovipositor clearly indicate this subfamily. The species was originally placed in *Pimpla*, but that was nearly fifty years before Henry Townes profoundly revised the generic classification of Darwin wasps (Townes 1969). While generic placement is ambiguous based on the holotype alone, the additional fossils here-in associated with this species are much better preserved. The yellow or orange ground colour and robust ovipositor point to either *Xanthopimpla* or to the *Theronia* group of genera. The in some specimens well-visible mandibles with two teeth exclude the former, which has twisted mandibles that appear unidentate in lateral or front view. Within the *Theronia* group, only *Epitheronia* has the lower tooth of the mandible distinctly shorter than the upper tooth.

*Species diagnosis*. The tri-coloured antenna distinguishes this species from all extant *Epitheronia*. There are two fossil species currently associated with the *Theronia* group, both described from the Miocene Florissant formation by Cockerell (1919): *Theronia wickhami* Cockerell and *Mesopimpla seqoiarum* Cockerell. We obtained photographs of the former from the University of Colorado Museum of Natural History, and it belongs to the Cremastinae. The whereabouts of the latter is unknown, but according to the original description, it is mainly black. Furthermore, drawings of the wing veins show that forewing vein 2m-cu is rather straight (bowed outwards in the new species) and that hindwing vein 1-Cu is longer than cu-a (distinctly shorter in *E. stigmatica*).

**Description.**

*Preservation*. Holotype rather poorly preserved, dorsal view. Head only partial, antennae missing except for some barely visible basal segments. Mesosoma poorly preserved, mesoscutum apparently broken off except for narrow, dark borders; left fore and hind wing well-preserved, traces of about two legs. Metasoma slightly better preserved, most tergites clearly or at least vaguely outlined, dark colouration only indicated in parts as slightly darker brown; ovipositor sheaths and base well preserved.

Best-preserved specimen (#MM-3141) shown in lateral view. Head including two complete antennae, mesosoma without much detail, forewings and a partial hind wing and large parts of all six legs present. Metasoma almost complete; ovipositor sheaths well preserved. Additional specimens either in ventral (#10099), dorsal (#11610) or lateral view (#11110, #11922), in two cases (#10099, #11922) with nearly complete antennae and in two with well-preserved legs showing enlarged claws (#11110, #11922).

Body 9.1–13.2 (9.1) mm. Yellow to light orange, with dark brown compound eyes and maybe head and marks on mesosoma, especially around wing base. Antennae orange on basal third, then with broad white ring, dark brown on apical 0.4. Wing veins dark except for light mark on basal ~3^rd^ of pterostigma; legs orange, hind tibia apically and hind tarsi darkened (see #11110). Dark markings at base of tergites 1 – at least 5 (see also discussion of colour preservation); ovipositor sheaths dark.

*Head* very short, with large compound eyes; in specimens #MM-3141 and #11922 with bidentate mandibles well visible, lower tooth distinctly shorter and a bit more narrow than upper tooth. Antenna about 1.1 times longer than fore wing, of even width, with about 34 to 39 flagellomeres; scape a little longer than wide, pedicel short, first flagellomere about 3.5 times longer than wide, subsequent ones decreasing in length to nearly quadrate in most apical flagellomeres.

*Mesosoma* rather short and stout; deep notauli converging on basal half, then nearly parallel (#11610); dark patches at fore wing base probably corresponding to axial sclerites; scutellum with transverse carinulae in prescutellar groove; propodeum higher than long, carination somewhat unclear, but seemingly with at least some closed areas and pleural carinae (#MM-3141, #11110). *Fore wing* 8.0–9.5 (8.0) mm; areolet closed, quadrate, with uneven sides, 4-M very short, 2r-m a bit shorter than 3r-m; 2m-cu curved outwards, with two bullae; pterostigma 3.5–4.2 (4.0) times longer than wide; 1cu-a meeting M + Cu a little posterior to 1-M; 3-Cu much longer than 2cu-a; radial cell 3.3–3.7 (3.3) times longer than wide. *Hind wing w*ith 1-Rs about 1.5 times longer than 1rs-m (holotype, #11610), 1-Cu 0.6–0.7 (0.65) times as long as cu-a. *Legs* rather well preserved, with two spurs at mid and hind tibial apex; hind femur about 2.9–3.6 (3.6) times longer than wide; claws rather large.

*Metasoma* rather stout, T1 about 0.8–1.0 (1.0) times as long as wide, with strong median longitudinal carinae converging on basal half (#11110); T2 0.5–0.7 (0.7) times as long as wide, T6 about the same length as T7, T8 very short. Ovipositor sheaths robust, 0.25 – 0.3 (0.3) times as long as fore wing; rather broad, covered with very dense and short hairs (#MM-3141).

*Theronia* Holmgren, 1859

*Theronia? furensis* **n. sp.**

Figure 8

**Figure 8.**
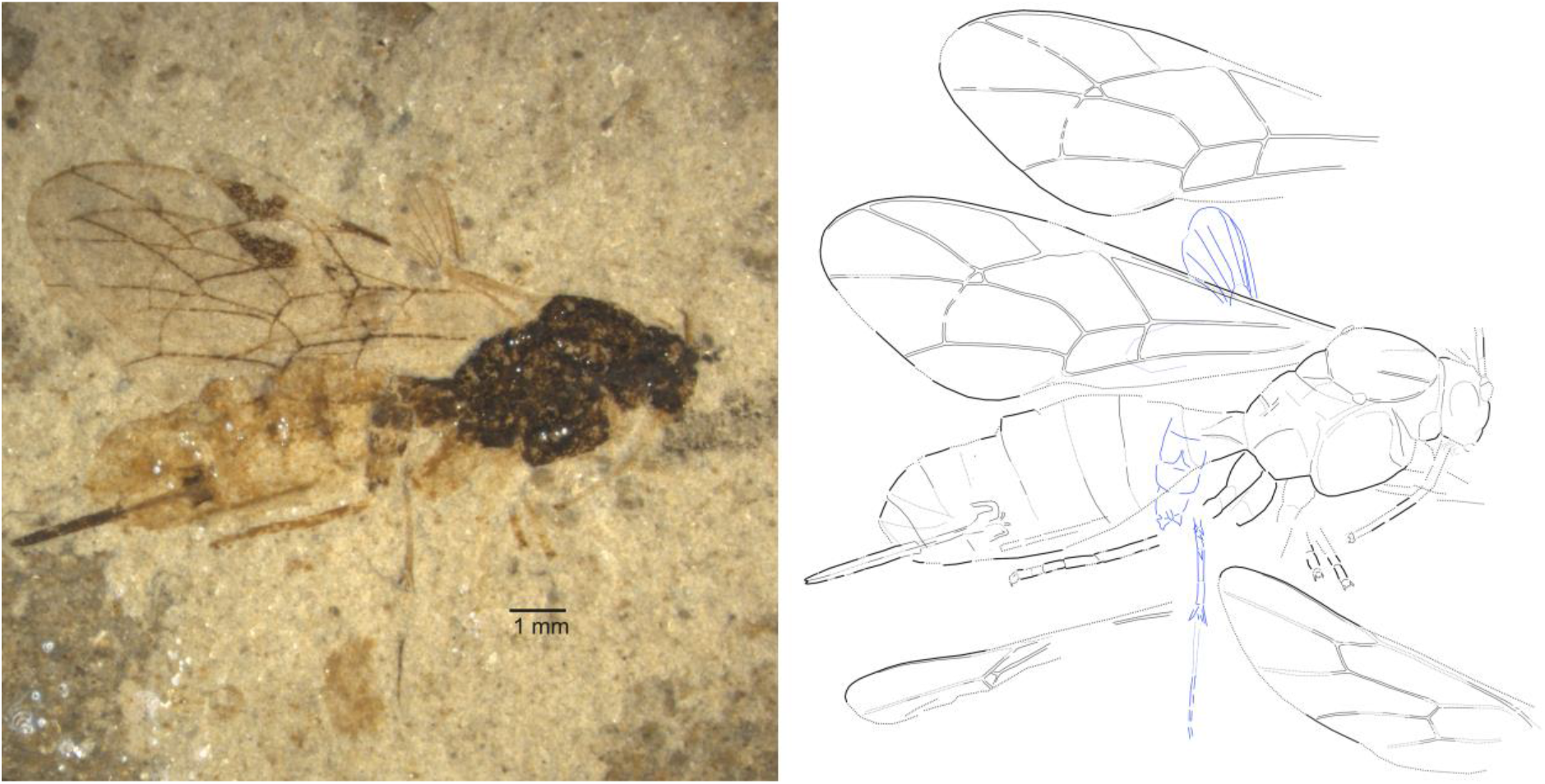
Holotype of *Theronia? furensis* (#11207), photograph and interpretative drawing, where dotted lines represent uncertain and/or interpolated interpretations. Blue lines indicate a fossil Diptera which is partly overlaying the Darwin wasp fossil. One fore and the two hind wings have been moved up and down, respectively, for clarity.

**Type material**. Holotype female: #11207 (part and counterpart; FUR; leg. E. Rettig).

**Etymology**. The species epithet refers to the formation of origin.

**Type horizon and locality**. Denmark, Jutland, Limfjord region, Morsø Kommune, Klinten ved Klitgård. Cement stone.

**Diagnosis.**

*Taxonomic placement*. The placement of this taxon in Pimplinae is rather unequivocal from the combination of characters indicated in Table 1. Furthermore, the claws seem somewhat enlarged, even though this interpretation is somewhat uncertain. The propodeum has at least some distinct carinae, a feature that is rare in extant Pimplinae which mostly have their propodeal carination at least partially reduced. The exception are some members of the genera *Crusopimpla, Xanthopimpla, Lissopimpla* Kriechbaumer and the *Theronia* group of genera, which all share a rather short metasoma and robust ovipositor with the current species. The first is a now extinct genus currently comprising only two species; we here added two additional ones and complemented the genus definition by an important character, which had not been visible in the previously known fossils: the high ratio of vein 1-Cu to cu-a in the hind wing (nervellus broken below the middle, see under *Crusopimpla violina*). *Crusopimpla* thus can be ruled out as placement for this fossil, which has vein 1-Cu a little shorter than cu-a. The remaining three genera would all fit in that respect, but the colour pattern of this species points to the *Theronia* group as a most likely placement: *Xanthopimpla* species have a light ground colour of the mesosoma, and *Lissopimpla* can be identified by extended light colouring on the mesosoma and a black patch in the fore wing. Placement in *Theronia* or any of its related genera must however be regarded as tentative, as most members of this genus group also have a yellow or orange ground colour of the mesosoma.

*Species diagnosis*. From most extant members of the *Theronia* genus group and also from most other pimplines from the Fur formation, this species can be distinguished on the basis of the colouration, i.e., dark mesosoma and head and entirely orange metasoma (except for T1) and orange legs. It only shares this colour pattern with *Crusopimpla? violina*, see under that latter species for differential characters.

**Description.**

*Preservation*. Lateral to dorso-lateral view. Head with first few segments of antennae. Mesosoma rather well preserved, including partial propodeal carination; all four wings nearly complete, all legs partially visible, often including claws. Metasoma almost complete; ovipositor sheaths nearly entirely preserved, with tip distinct. Partial Diptera fossil with wing fragment, partial legs and partial abdomen overlaying wasp fossil.

Body 11.1 mm. Black or dark brown, orange on mouth parts, at least base of antennae, all legs including coxae and tarsi, metasoma from tergite 2, ovipositor sheaths dark brown; basal ~3^rd^ of pterostigma whitish.

*Head* rather small and roundish, antenna with short scape and pedicel.

*Mesosoma* rather short and stout; deep notauli visible over at least a third of mesoscutum, a little converging but still far apart when they become invisible; scutellum rather short; mesopleuron at least with lower and probably also upper lateral portion of epicnemical carina visible. Propodeum about as long as high, with some distinct carination, corresponding to at least one transverse and a partial median longitudinal and pleural carinae. *Fore wing* 9.5 mm; areolet closed, quadrate, with uneven sides, 4-M very short, 2r-m a bit shorter than 3r-m; 2m-cu with two bullae, clearly bent outwards; pterostigma about 3.5 x longer than wide; 1cu-a meeting M + Cu slightly distal to 1-M; 3-Cu clearly longer than 2cu-a; radial cell 3.2 x longer than wide. *Hind wing w*ith 1-Rs 1.5 x longer than 1rs-m, 1-Cu a little shorter than cu-a (counterpart), but end of the latter a bit uncertain. *Legs* all partially preserved, mostly with femora and tibiae rather indistinct, but tarsi visible even with claws, some of which vaguely show what is probably a basal lobe.

*Metasoma* rather stout, with T1 less than 1.5 x as long as wide, with strong median longitudinal carina converging at least on basal two-thirds; T2 transverse, less than 0.8 x as long as wide, remaining segments even shorter. Ovipositor sheaths ~0.25 x as long as fore wing, their tips partly open to allow extrusion of ovipositor tips.

*Theronia? nigriscutum* **n. sp.**

Figure 9

**Figure 9.**
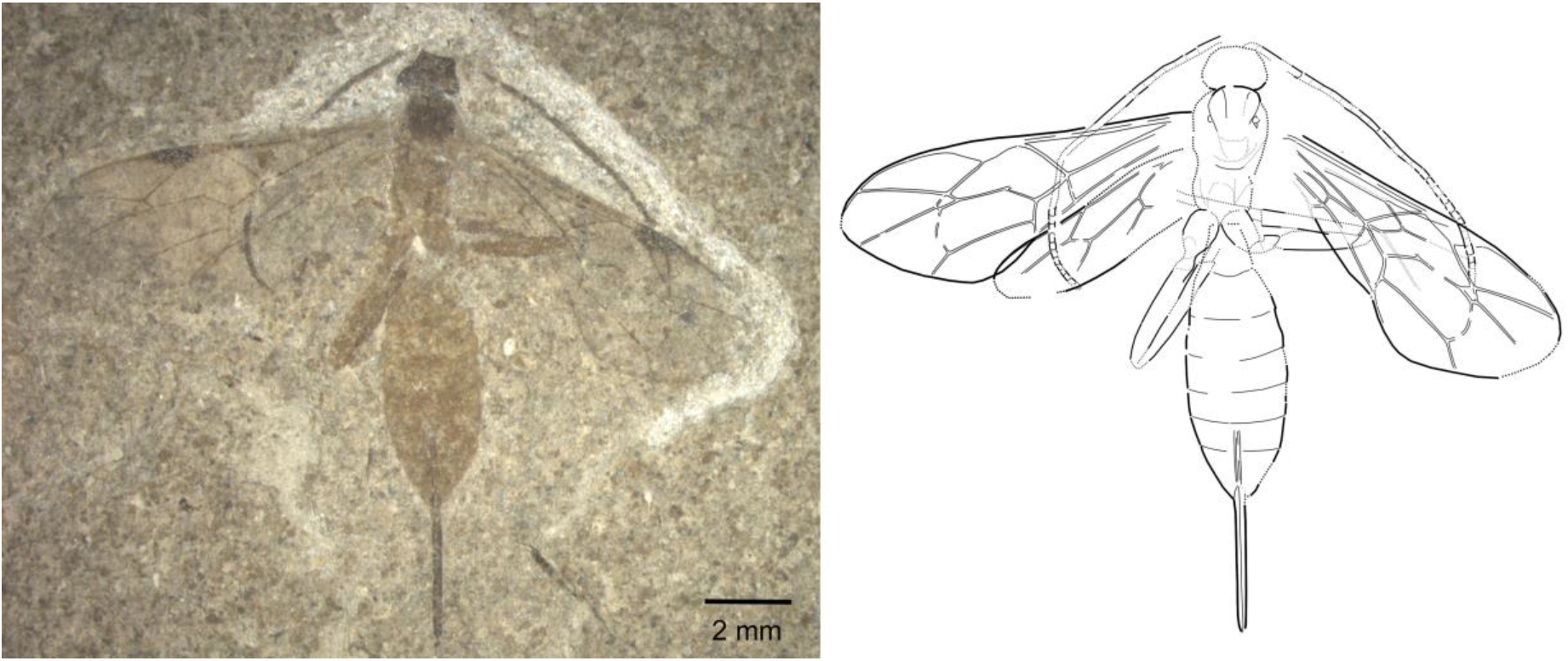
Holotype of *Theronia? nigriscutum* (#MHM 5412), photograph and interpretative drawing, where dotted lines represent uncertain and/or interpolated interpretations.

**Type material**. Holotype female: #MHM-5412 (part and counterpart; MOL; leg. H. Madsen, 1995).

**Etymology**. This species is named for its dark mesoscutum.

**Type horizon and locality**. Denmark, Jutland, Limfjord region, Morsø Kommune, Ejerslev. Silstrup Member, +25-30 (Blok “E”).

**Diagnosis.**

*Taxonomic placement*. This species shows most of the typical pimpline characters listed in Table 1, although the ovipositor appears somewhat more slender than in other species of this subfamily from the Fur Formation. The nervellus, which is broken above the middle, and the yellow or orange ground colour fits well with members of the *Theronia* genus group, although the current specimen appears somewhat more elongate, especially in terms of the hind femora. Generic placement must thus be regarded as tentative only, as many diagnostic characters such as details of the propodeum are not well visible in the single specimen of this species.

*Species diagnosis*. This species shares its colour pattern with some *Xanthopimpla* and *Theronia*-group species, including *X. crescendae* and *Epitheronia stigmatica* from the same formation. From the latter two, it can easily be distinguished by the uniformly dark colour of the antennae. It is in general more slender than all known extant and fossil *Xanthopimpla* and most *Theronia*-group species, especially in terms of the hind legs.

**Description.**

*Preservation*. Dorsal view, but T1 mostly replaced by hind coxae. Head large parts of both antennae present. Mesosoma with mesoscutum rather well preserved, remainder somewhat less so; front wings complete, hind wings almost complete, hind legs partially visible. Metasoma almost complete, except first tergite, in whose place the hind coxae show rather prominently; ovipositor sheaths nearly entirely preserved, with tip distinct.

Body 10.8 mm. Orange with dark brown head, antennae, mesoscutum, wing veins, and ovipositor sheaths.

*Head* rather small and roundish, antenna with about 30 flagellomeres, basal segments about 1.5 times longer than wide, median and apical segments square to transverse.

*Mesosoma* rather slender; deep notauli visible over at least half of mesoscutum, first a little converging, then nearly parallel. Propodeum not well preserved, maybe with some carination indicated. *Fore wing* 8.7 mm; areolet closed, quadrate, with uneven sides, 4-M very short, 2r-m a bit shorter than 3r-m; 2m-cu with two bullae, clearly bent outwards; pterostigma about 3.8 x longer than wide; 1cu-a meeting M + Cu opposite of 1-M; 3-Cu clearly longer than 2cu-a; radial cell 3 x longer than wide. *Hind wing w*ith 1-Rs 1.5 x longer than 1rs-m, 1-Cu probably about 0.8 times as long as cu-a, but end of the latter a bit uncertain. *Hind legs* partially preserved, rather slender, hind femur not conspicuously thickened as in other Fur Formation pimplines with yellow or orange ground colour.

*Metasoma* rather stout, with T1 a little longer than wide; T2 transverse, about 0.6 x as long as wide, remaining segments even shorter. Ovipositor sheaths ~0.4 x as long as fore wing.

*Xanthopimpla ciboisae* **n. sp.**

Figure 10

**Figure 10.**
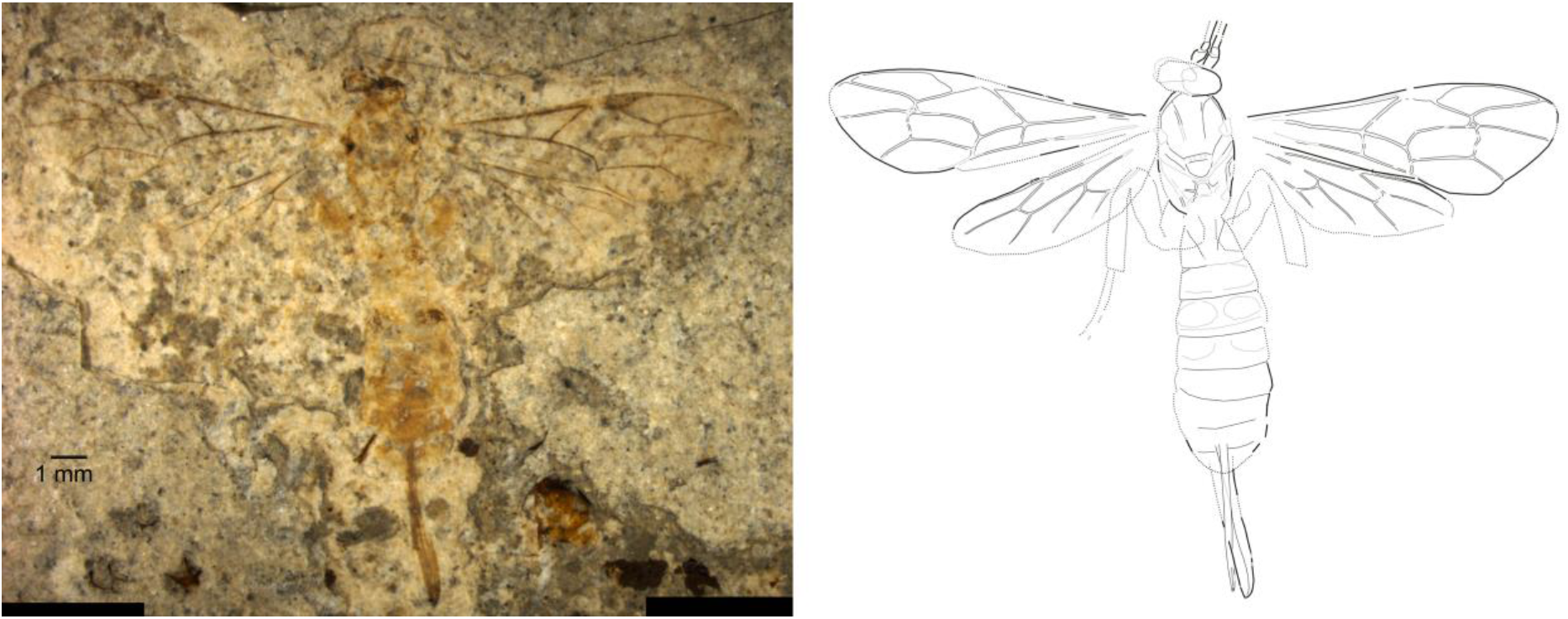
Holotype of *Xanthopimpla ciboisae* (#10046), photograph and interpretative drawing, where dotted lines represent uncertain and/or interpolated interpretations.

**Type material**. Holotype female: #10046 (FUR; leg. M. Breiner).

**Etymology**. This wasp is dedicated to Alice Cibois, to honour her dedication as president of the Swiss Systematics Society and her creative and innovative efforts to promote our craft among the general public. She especially drew attention to the ongoing discovery and naming of new species through the curated list “New Species Swiss Made”, which year after year feature dozens to hundred new species across the tree of life.

**Type horizon and locality**. Denmark, Jutland, Limfjord region, Skive Kommune, Fur. Silstrup member.

**Material examined**. Tentatively placed in this species: #14684 (FUR).

**Diagnosis.**

*Taxonomic placement*. While many of the characters that can be used to diagnose Pimplinae are well visible in this specimen (Table 1), the most unequivocal evidence stems from generic placement: an association with *Xanthopimpla* is evidenced by the colour pattern (especially the paired dark marks at least on tergite 3, robust legs, very short hind wing 1-Cu compared to cu-a, stout metasoma, and robust ovipositor. Forewing vein 2m-cu furthermore shows a zig-zag pattern that is very rare outside that genus.

*Species diagnosis*. This species is very similar both in colour pattern and wing venation to many extant species of the genus, but has a more extended light mark at the base of the pterostigma. It is much larger than the two fossil species from Messel, even than *X. biamosa* (forewing 8.0 mm), from which it can be further distinguished by the lateral dark marks on T1 in the latter. From *X. stigmatica*, it can be distinguished by the larger size, light coloured mesoscutum, more sinuous 4Rs in the front wing, and even shorter 1-Cu compared to cu-a in the hind wing.

**Description.**

*Preservation*. Dorsal view. Head partially preserved, including extreme base of antennae. Mesosoma rather well preserved, including propodeal carination; all four wings nearly complete, partial hind legs visible. Metasoma almost complete; ovipositor partly, sheaths nearly entirely preserved.

Body 11.3 mm. Yellow to light orange, with dark brown compound eyes, wing base, wing veins except basal ~3^rd^ of pterostigma; dark paired spots evident on T3, seemingly absent from the other tergites (see also discussion of colour preservation); ovipositor light orange, sheaths brownish.

*Head* distorted, with compound eyes overlapping; scape, pedicel and part of first flagellomeres preserved, of normal dimensions.

*Mesosoma* rather short and stout; deep notauli converging on basal ¼, then parallel; dark patches at fore wing base probably corresponding to axial sclerites; scutellum rather short, with transverse carinulae in prescutellar groove; propodeum with distinct carination, corresponding to nearly complete basal and probably also apical transverse, lateromedian and maybe lateral longitudinal and pleural carinae. *Fore wing* 9.2 mm; areolet closed, slightly petiolate above, quadrate with uneven sides (4-M very short, 2r-m a bit shorter than 3r-m); 2m-cu with two bullae, with a weak zig-zag that anteriorly is bent outwards, then inwards in front of the posterior bulla; pterostigma 3.5 x longer than wide; 1cu-a meeting M + Cu opposite 1-M; 3-Cu a bit longer than 2cu-a; radial cell 3.5 x longer than wide. *Hind wing w*ith 1-Rs about 3 x longer than 1rs-m, 1-Cu 0.3 x as long as cu-a. *Hind leg* not well preserved, but rather stout.

*Metasoma* with T1 about 0.7 x longer than wide, with strong median longitudinal carina converging at least on basal two-thirds; T2 0.4 x times as long as wide, with basal oblique grooves indicated and maybe a preapical transverse groove present. T3 with paired dark patches on basal half and pre-apical transverse groove indicated. T4 with weak shadows of paired marks, unclear on remainder of tergites. Ovipositor sheaths ~0.35 x as long as fore wing; rather broad, covered with very dense and short hairs.

*Xanthopimpla* Saussure, 1892

*Xanthopimpla crescendae* **n. sp.**

Figure 11

**Figure 11.**
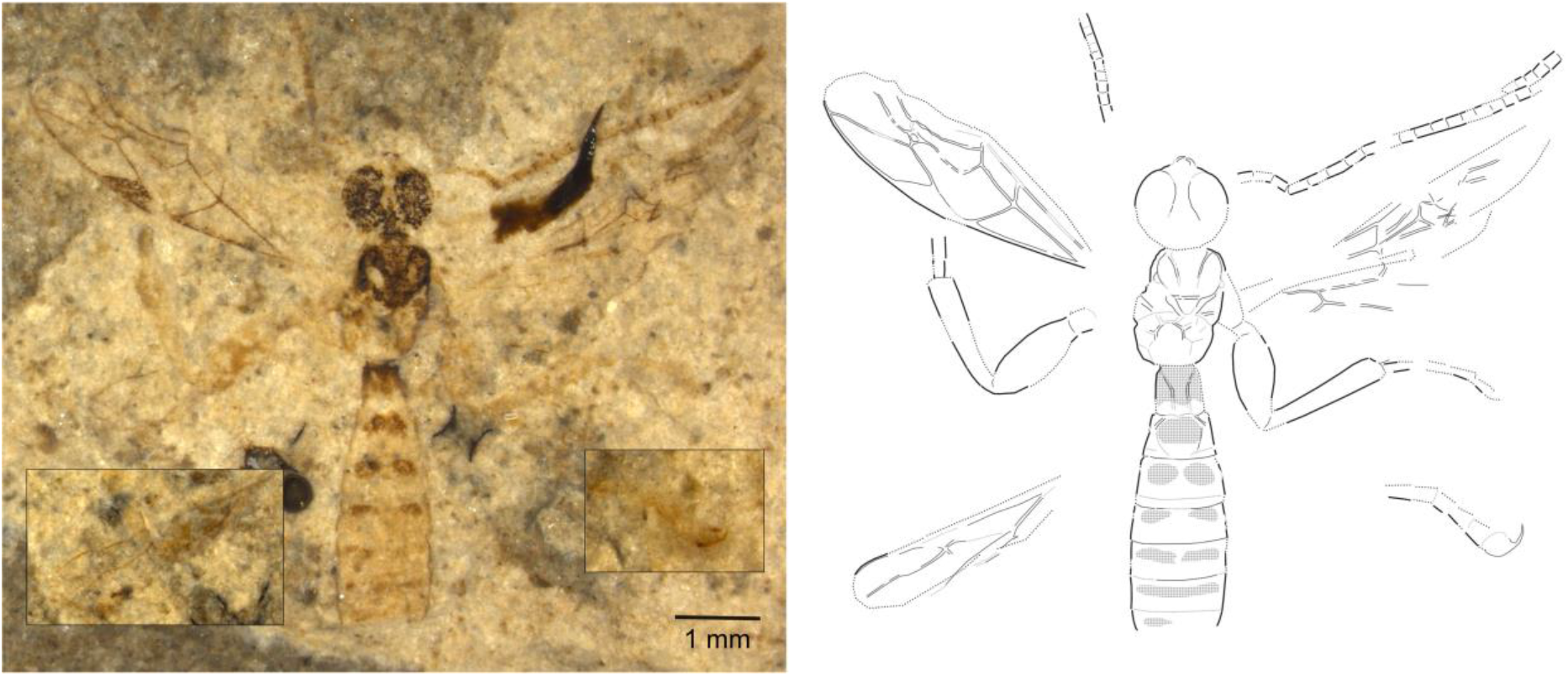
Holotype of *Xanthopimpla crescendae* (#11214), photograph and interpretative drawing, where dotted lines represent uncertain and/or interpolated interpretations.

**Type material**. Holotype #11214 (male?, part and counterpart; FUR; leg. E. Rettig), paratypes #13065 (FUR; female), #10918 (FUR; sex unknown).

**Etymology**. This wasp is dedicated to my dear friend Tabia Stoffel, nickname “Crescendo”, for her unwavering support over the years.

**Type horizon and locality**. Denmark, Jutland, Limfjord region, Morsø Kommune, Klinten ved Klitgård (holotype); Svalklit (#13065). Skive Kommune, Østklinten (#10918). Cement stone.

**Material examined**. Tentatively identified as the same species: #13068, #16642, #17232, #17235 (all FUR).

**Diagnosis.**

*Taxonomic placement*. The areolet shape, outwards-bowed 2m-cu in the forewing, as well as the short and stout T1 and robust, protruding ovipositor clearly indicate Pimplinae. But most of all, there is clear evidence for placing this species in *Xanthopimpla*: narrow (twisted) mandibles (visible in holotype), yellow or orange ground colouration with paired, dark markings on several tergites, stout legs, extensive carination of the propodeum, hind wing with 1-Cu shorter than cu-a (nervellus broken above middle), large claws.

*Species diagnosis*. This species stands out within the genus by its almost or entirely dark mesoscutum and broad light mark at the base of the pterostigma, a combination not seen in any extant nor in the three known fossil species of the genus. *Xanthopimpla biamosa* Khalaim, 2008 from the Oligocene of Biamo (Russia) furthermore has much shorter and stouter hind femora and two lateral dark marks on tergite 1, instead of its base being dark. *X. messelensis* Spasojevic et al., 2018 from the mid Eocene deposit in Messel has a very narrow, triangular areolet and dark basal bands instead of clearly disconnected spots on tergites 2-3. And *X. praeclara* Spasojevic et al. 2018, also from Messel, can be distinguished by the entirely yellow tergite 2. Among the here-in described Fur fossil species, *X. nigriscutum* is most similar to *X. ciboisae*, from which it can be distinguished by its dark mesoscutum, smaller size and more slender metasoma.

**Description.**

*Preservation*.

Holotype preserved in dorsal view, with head including most of one and a partial second antenna, most of mesosoma, one nearly complete but somewhat distorted and one longitudinally folded forewing, two partial hind wings, partial hind legs, and metasoma except for its tip. Metasoma especially well-preserved, including traces of microsculpture on the tergites. Paratype in dorsal view, head without antennae, most of mesosoma, partial fore wings, one nearly complete hind wing, metasoma except for its tip.

Body ~7.8– 9.3 (~7.8) mm. Yellow to orange, with dark brown head, antennae uniformly orange. Dark on entire mesoscutum except maybe two lateral patches, basal part of propodeum, wing veins except basal half of pterostigma, basal two thirds of T1, paired spots on T2 to T4, on T5-6 these spots combining into a basal dark stripe.

*Head* in holotype turned upward to show from front, rather high with large compound eyes, these seemingly converging towards ventral (but this might be an artefact); with very narrow, darkened and bent mandibular tips than might indicate either a single tooth or strongly narrowed and twisted mandibles. Antennae incomplete, showing at least 23 clearly outlined flagellomeres (estimated: 26 in total), these in proximal half about twice as long as wide, after middle about 1.5 times, and penultimate segments about as long as wide.

*Mesosoma* rather slender, in holotype distorted between mesoscutum and propodeum. Mesoscutum about as long as wide, with strongly converging but not meeting notauli. Propodeum with carination partially preserved (holotype), showing at least basal transverse, and lateral parts of apical transverse, and part of the median longitudinal carinae. *Fore wing* 5.5–6.3 (5.5) mm; areolet closed, quadrate, with uneven sides, 4-M very short, 2r-m a bit shorter than 3r-m; 2m-cu not well preserved in holotype but clearly bowing outwards in paratype; pterostigma 3.7–3.9 (3.7) x longer than wide; 1cu-a meeting M + Cu opposite 1-M; 3-Cu a bit longer than 2cu-a; radial cell 3.2–3.3x (3.3) longer than wide. *Hind wing w*ith 1-Rs about 1.8 x (paratype) longer than 1rs-m, 1-Cu 0.5-0.7 x (0.5) as long as cu-a. *Hind leg* very short and stout, femur about 2.5x longer than wide. Tarsal claw discernible, rather large.

*Metasoma* with T1 about 1.1x longer than wide, with strong median longitudinal carina converging on basal half, parallel posteriorly; T2 0.6 times as long as wide, basally with oblique grooves cutting off anterolateral corners, behind these with slowly converging grooves that cut off a roundish-trapezoid raised part medially bearing two confluent dark patches, this part with strong punctures visible. T3 with paired dark patches on basal half and pre-apical transverse groove indicated. T4-T7 with paired dark marks weakly indicated. Ovipositor not preserved.

## Discussion

### How reliable are colour characters found in Fur Formation fossils?

We found colouration in the studied fossils very useful both to delimit species and to inform their placement in extant genera. But is colour preservation in these fossils sufficiently reliable to serve such taxonomic purposes? Colour can be strongly during diagenesis, and the outcome not only depends on the type and distribution of colouration present in the organism, but also on various aspects of the fossilization process, such as the biological and chemical composition of the sediment and the regime of temperature and pressure that the specimen gets exposed to (Mcnamara 2013). This is the case both for structural colours, which can change their hue drastically without showing much distortion in the colour-producing ultrastructure (Cai et al. 2020, Mcnamara et al. 2011), and for colour that results from pigments.

On the other hand, evidence is currently accumulating that not all colouration might be lost during fossilization. Structural colouration can be reconstructed from the biophotonic nanostructures, if these are sufficiently well preserved (Mcnamara 2013). And it has been shown convincingly that at least melanins, the most common class of pigments in insects, might be preserved nearly unaltered under certain fossilization conditions, for instance in the ink bladder of a Jurassic squid (Glass *et al*. 2012) and in melanosomes in a fish eye from the Fur Formation (Lindgren et al. 2012). However, these specimens might represent exceptional conditions, and it remains unclear whether such pigment preservation is also possible in insect cuticle. Nevertheless, the process might still preserve some of the original patterns, if not the hues of the original colouration. The origin of the sometimes very detailed patterning in different shades of brown in many insect fossils, including those studied here, is still poorly understood, but besides original pigmentation, it might also reflect variations in the general composition of the cuticle, which in turn might be correlated to original colour patterns. This is because melanins are not only involved in signalling or camouflage, but also in the sclerotization of the cuticle (Andersen 2010).

The parallels in colour patterns especially in the *Xanthopimpla* species described here-in with extant representatives of the genus are rather striking. This observation is supported further by three other fossil *Xanthopimpla* species, two from the Eocene Messel pit in Germany (Spasojevic *et al*. 2018b) and one from the Oligocene Biamo Formation in Russia (Khalaim 2008). While the respective hues are somewhat different between the localities, type and locations of dark-and-light patterns on the body are consistent with each other and with extant members of the genus. Fossilization conditions must have been rather different among these three fossil localities now known to harbour species of this genus: Messel was a largely anoxic crater lake with fossils enclosed in oil shale, Biamo represents a lacustrine environment with diatomite sediments, while Fur consists of marine sediments.

In brief, although it is not very likely that the original pigments cause the colours observed today in the studied fossils, comparisons among fossil and between them and extant taxa suggest that they at least in part reflect original colour patterns, although the precise mechanisms that preserved them for dozens of millions of years remain to be unravelled. The difference in the consistency of colour preservation between dark-bodies and light-bodied ichneumonids from the Fur Formation deserves further attention, for instance by examining differences in the cuticular ultrastructure in extant and fossil representatives with both dark and light ground colour of the body.

### Diversity of Pimplinae in Fur Formation fossils

The Fur Formation pimplines are the oldest representatives of this subfamily known so far, although this status might be challenged soon by some species currently placed in Labenopimplinae that might turn out as pimplines, such as *Rugopimpla* (Dmitry Kopylov, personal communication). The ten species of pimpline wasps covered here suggest a high diversity of this subfamily in the Fur Formation, which is otherwise only mirrored by the late Eocene Florissant Formation (Mitchell 2013-although many of the species described from the latter are in need of revision and might turn out not to be pimplines after all). From the early Eocene, three species have been described from the Green River Formation (Spasojevic et al. 2018a) and four from the Messel pit (Spasojevic et al. 2018b). If only considering named species from the fossil record, pimpline diversity in Fur would seem extraordinary; however, it is actually quite low when compared to species numbers of this subfamily in many extant terrestrial habitats, which can easily amount to several dozens or hundreds. Potential reasons for this discrepancy are manyfold. First, behavioural and phenological differences between species might have made some more prone to being blown out by storms into the ocean. Second, additional specimens of Darwin wasps are excavated at a regular pace in the Fur Formation, and many additional species can thus be expected in the coming years. Third, our approach to species delimitation was rather conservative, requiring strong evidence for the erection of additional taxa. In extant Darwin wasps, differences between species often manifest in details in the colouration or even microsculpture of specific parts of the cuticle (e.g., Klopfstein 2014, Pham et al. 2011) – characters not easily observed in fossils. We have thus certainly underestimated species numbers in Fur Formation pimplines.

The diversity within Pimplinae appears skewed towards particular genera and body shapes when compared to extant communities. A recent study using total-evidence dating to calibrate the phylogeny of Pimpliformes (Spasojevic et al. 2021) indicates that all the pimpline tribes started radiating already deep in the Cretaceous, suggesting that another explanation than age is needed to resolve this discrepancy. This remains true even when considering that Pimplinae as the subfamily is currently understood is probably paraphyletic (Klopfstein et al. 2019a). Instead, behavioural differences might play a role here, but the pattern might also indicate different habitat requirements. For instance, all ovipositors that were well-preserved among the studied pimplines had about the same relative length, between 0.2 and 0.4 times the forewing. However, there is a rich diversity of extant species with much longer ovipositors, for instance in the pimpline tribe Ephialtini. They typically attack hosts deeply concealed in wood, such as wood wasp or coleopteran larvae. Any such species are currently missing from the Fur Formation assemblage, but are present and even prevalent in other localities, such as the Florissant Formation. If this pattern is confirmed, it might indicate that the habitat of origin of the Fur Formation insects was rather open land than dense forest. However, a more detailed study of the exquisite insect fossil record from the Fur Formation, ideally including detailed information about the exact horizon, is needed to draw firm conclusions about ecological requirements of the respective insect communities.

## Disclaimer

This preprint has been submitted for publication in a peer-reviewed journal. All original descriptions in this bioRxiv document are not issued for public and permanent scientific record, or for purposes of zoological nomenclature. They should only become available with the proper publication of the article.

## Acknowledgment

I am indebted to René Lyng Sylvestern (Salling Museum in Fur) who kindly provided me with some 30 kilograms of Fur Formation fossils. Lars Vilhelmsen (Natural History Museum of Copenhagen) provided access to the holotype of *Pimpla stigmatica* Henriksen and assisted by taking measurements. I am grateful to all the collectors who tirelessly and carefully excavate hundreds of insect fossils and make them available to science by donating them to the local museums. Alexandra Viertler (Natural History Museum in Basel) identified some of the specimens, and Bastien Mennecart provided valuable feedback on the manuscript and translations into French. Dmitry Kopylov made important suggestions that helped improve considerably a previous version of this manuscript. This work was supported by grant PZ00P3_154791 of the Swiss National Science Foundation.

